# The immunopeptidome from a genomic perspective: Establishing immune-relevant regions for cancer vaccine design

**DOI:** 10.1101/2022.01.13.475872

**Authors:** Georges Bedran, Tongjie Wang, Dominika Pankanin, Kenneth Weke, Alexander Laird, Christophe Battail, Fabio Massimo Zanzotto, Catia Pesquita, Håkan Axelson, Ajitha Rajan, David J. Harrison, Aleksander Palkowski, Maciej Pawlik, Maciej Parys, Robert O’Neill, Paul M. Brennan, Stefan Symeonides, David R. Goodlett, Kevin Litchfield, Robin Fahraeus, Ted R. Hupp, Sachin Kote, Javier A. Alfaro

## Abstract

A longstanding disconnect between the growing number of MHC Class I immunopeptidomic studies and genomic medicine hinders cancer vaccine design. We develop COD-dipp to genomically map the full spectrum of detected canonical and non-canonical (non-exonic) MHC Class I antigens from 26 cancer studies. We demonstrate that patient mutations in regions overlapping physically identified antigens better predict immunotherapy response when compared to neoantigen predictions. We suggest a vaccine design approach using 140,966 highly immune-visible regions of the genome annotated by their expression and haplotype frequency in the human population. These regions tend to be highly conserved, mutated in cancer and harbor 7.8 times more immunogenicity. Intersecting pan-cancer mutations with these immune surveilled regions revealed a potential to create off-the-shelf multi-epitope vaccines against public neoantigens. Here we release COD-dipp, a cancer vaccine toolkit as a web-application (https://www.proteogenomics.ca/COD-dipp) and open-source high-throughput resource.

## Introduction

A revolution in cancer vaccines is underway fueled by a better understanding of the immune response and the breadth of tumor-associated antigens (neoantigens)^1^. In part, this is due to the accelerated adoption of MHC Class I antigen profiling (*i*.*e*., immunopeptidomics) in cancer by mass spectrometry. Leveraging these studies for cancer vaccines involves connecting the wealth of immunopeptidomics data to immunogenomics, where the goal is to carefully choose effective mutations to develop vaccines^2^.

The immunopeptidomics community has emphasized linking genomics to proteomics through the direct detection of neoantigens among the unexplained spectra in mass spectrometry studies^3–5^. Yet, there are known sequence coverage limitations of current MS-based proteomics that limit the detection of neoantigens and therefore the effectiveness of this approach^6,7^. Oppositely, genomics has emphasized the use of MHC binding predictors^8– 10^ trained from immunopeptidomics and affinity data to propose neoantigens for vaccines. Although their artificial intelligence architecture offers the most predictive accuracy, they still suffer from some drawbacks. First, the training datasets lag behind on cutting edge developments in computational mass-spectrometry that would identify non-canonical antigens. Second, they do not generalize well to the highly polymorphic MHC alleles^11^ or to peptides with different lengths. Third, a high fraction of peptides is predicted to bind strongly, where in fact they do not, meaning a high proportion of these predicted neoantigens will not be effective for vaccine design^12^. Finally, an accurate landscape of the presented immunopeptidome remains inaccessible within these predictive models, breaking the flow between the detected genomic aberrations and their predicted MHC presentation.

From a therapeutic standpoint, cancer vaccines have been designed to target private neoantigens^13^ personalized to individual patients’ tumors. This strategy presents a substantial bottleneck in terms of production and scalability because vaccines must be tailored to each patient. A more sustainable strategy is to use public neoantigens^14^ that rely on recurrent mutations in cancer to develop antibody and T cell therapies. Thus, a comprehensive list of such recurring neoantigens broadly presented to the immune system across the human population is urgently needed. These alternative therapeutic agents could cover the genomic diversity of presented antigens as well as the multitude of co-translational^15^ and post-translational^16^ aberrations. Such ‘focal public neoantigens’ could be used to refine the development of multi-epitope vaccines against cancer and could offer a new line of population-level immunotherapeutics.

Since cancer is a disease of the genome, tracing the physically detected immunopeptidome back to the genome stands to transform cancer vaccine design, yet remains difficult. To start, no high-throughput method for capturing the full-breadth of the immunopeptidome including canonical (exonic and post translationally modified) and non-canonical (intronic, frame shifted or UTR peptides) has been put forward. In addition, harmonized analyses have yet to connect accessibly to the genome-centric bioinformatics community.

Here, we present a comprehensive and well-engineered resource of presented peptidic antigens (immunopeptidomes) that incorporates the full breadth of neoantigen science^17–21^. The immunopeptidome profiles of over 486 samples from 26 published cancer studies^22,23^ were analyzed using a novel harmonized approach. Assembling a novel catalog of peptides derived from canonical (exonic or post translationally modified) and non-canonical (intronic, frame shifted or UTR) sources uncovered a spectrum of recurrent in-frame antigens and out-of-frame neoantigens. The genome centric nature of our resource makes the connection to cancer mutation data simple. Aligning these peptides to the genome revealed a cartography of over 468,048 unique immunopeptides and proved that mutations in the corresponding regions are predictive for immunotherapy response. Our pan-cancer analysis relying on focal public neoantigens suggests which multi-epitope vaccines will make strong candidates for the next generation of vaccines and T-cell based therapies. We assess the feasibility of designing poly-epitope vaccines against focal public neoantigens and provide data, alongside a web-application, to explore and develop these vaccines for broad spectrum therapeutics. Beyond cancer, the methods introduced here stand to enable important questions about the status and evolution of the immune-visible genome for the comparative immunology community.

## Results

### Immunopeptidomics Mass Spectrometry datasets

We selected 26 immunopeptidomics mass spectrometry studies **(Supplementary Table 1)** to create our dataset of antigen presentation in cancer. Samples were analyzed using a harmonized approach for the characterization of data-dependent acquisition (DDA) mode in tandem mass spectrometry. These DDA datasets covered several cancer types affecting brain, lung, skin, liver, blood, colon, ovarian, and breast cancer tissues, (**Fig. 1a, methods, Supplementary Note 1**) including cell lines, and disease free normals (**Fig. 1b**). We filtered the available data for high-resolution mass spectrometry instruments (Q Exactive Plus/HF/HFX and Fusion Lumos) to minimize the bias associated with older tandem mass-spectrometry instrumentation (**Fig. 1c**). Antibody choice can impact which MHC molecules are selected for analysis in immunoprecipitation (IP)^24^. Within our dataset, W6/32 was the most used monoclonal antibody for HLA class I IP compared to the other antibodies (BB7.2 and G46-2.6) (**Fig. 1d**, cf. **Supplementary Table 1**). The studies cover 5 different HLA Class I genes with HLA-A, B, and C being the most studied compared to HLA-E, and G (**Fig. 1e**).

**Figure 1:**
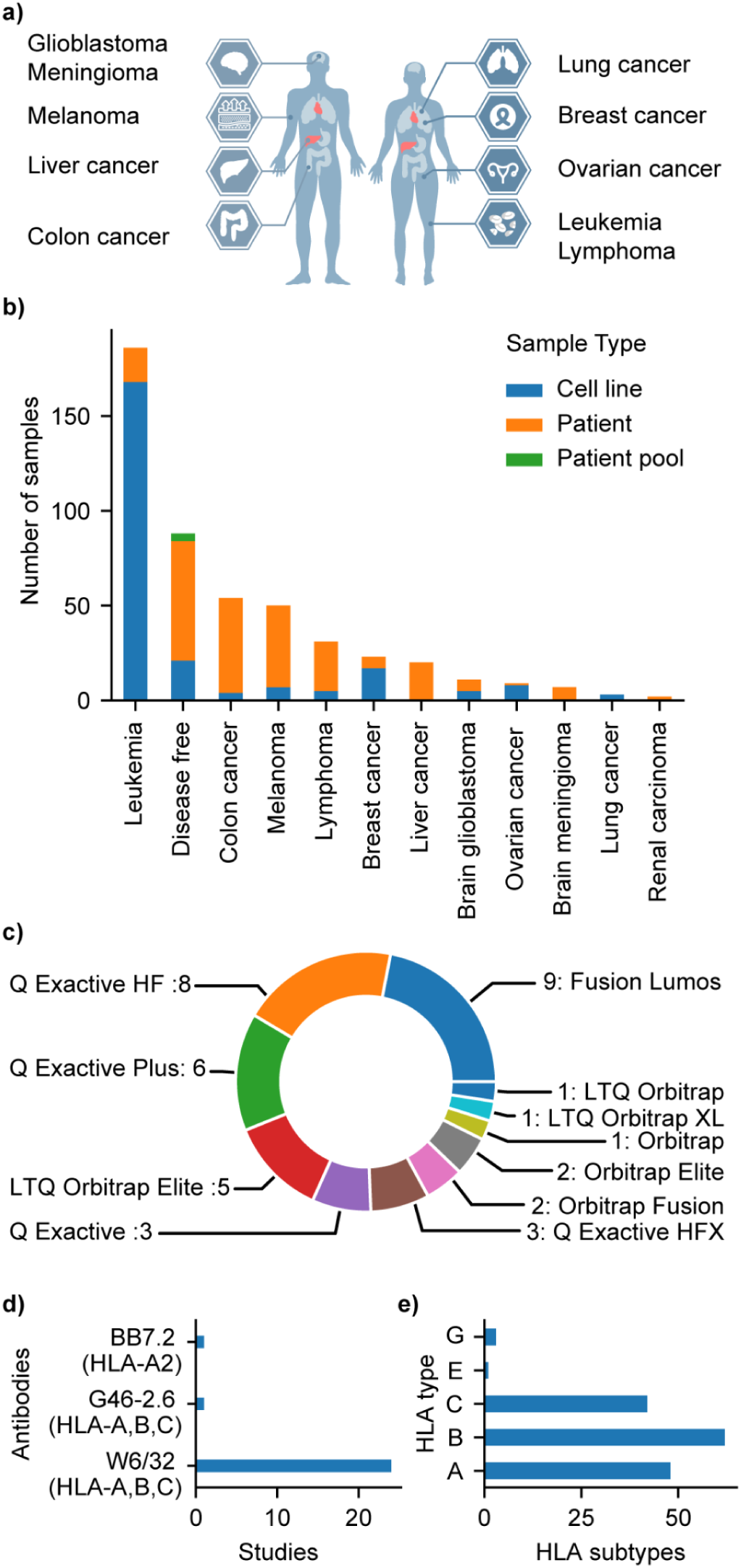
Included immunopeptidomic datasets. **(a)** Different types of cancers considered in this study. **(b)** Number of cell lines (blue) patient tissues (orange) and patient pools (green) per lesion type **(c)** Mass spectrometry instruments considered (instrument name: number of studies) **(d)** Antibodies used for Immuno-precipitation (IP) **(e)** Overall number HLA subtypes per HLA type

### COD-dipp: A high-throughput pipeline for the interrogation of immunopeptidomics datasets

We present COD-dipp (Closed Open Denovo - deep immunopeptidomics pipeline), an open-source high throughput pipeline with novel post-processing steps to deeply interrogate immunopeptidomics datasets (**Fig. 2a)**. We then use this pipeline to develop a well-engineered database of canonical and non-canonical MHC Class I peptides, and an accessible web-application to facilitate their use for vaccine design (**Fig. 2b**). We chose to work with DDA datasets, owing to their abundance in the literature. DDA data can be analyzed using different computational methods to identify the peptides by matching the acquired MS2 spectra to an amino-acid sequence^25^ in a process called peptide-spectrum matching (PSM)^26^. Closed search, open search, and *de novo* sequencing are three main strategies used to identify canonical, post-translationally modified and non-canonical peptides respectively. We chose one algorithm from each of these categories of PSM assignment: MS-GF+^27^, MSFragger^28^, and deepNovo v2^20^ in order to cover more of the spectra in all 486 samples. We interfaced the deep neural network (deepNovoV2) with the closed search algorithm (MS-GF+) to automatically learn the interpretation of mass spectrometry spectra. Using our novel *de novo* post-processing we traced these peptides to canonical and non-canonical sources using carefully tuned short sequence alignments (**Supplementary Note 2; Supplementary Fig. S1**).

**Figure 2:**
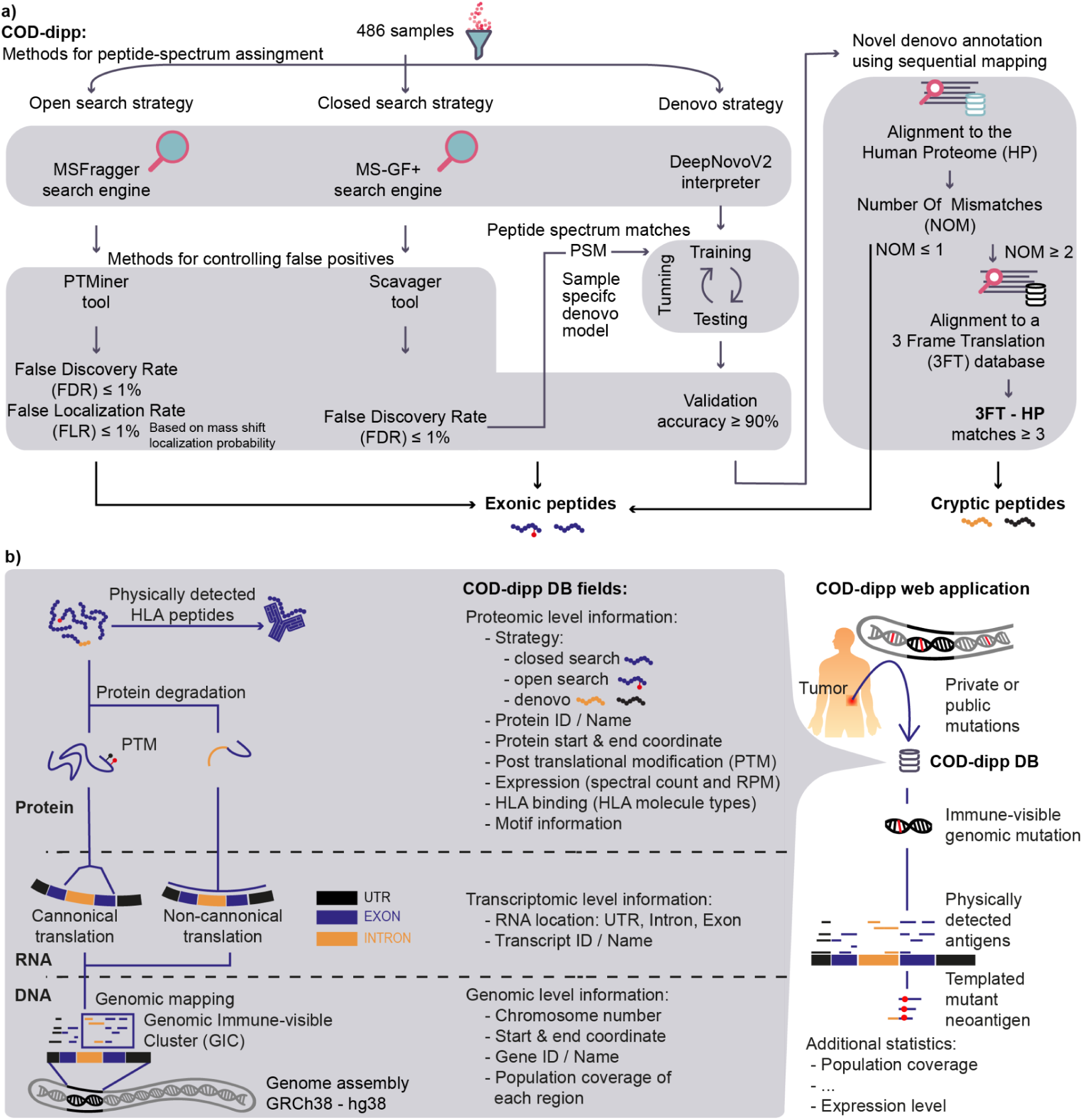
COD-dipp: A new high-throughput pipeline for a deep interrogation of immunopeptidomic datasets. **(a)** Three strategies of peptide spectrum assignment were combined: closed search (for canonical peptides), open search (for PTMs), and de novo (for canonical and non-canonical peptides). False Localization Rates for PTMs and False Discovery Rates for peptide calling were carefully controlled to 1%. An approach to find non-canonical peptides developed here uses a denovo sequencing model trained for each sample using the quality controlled peptide-spectrum matches from closed search. Results were split into 3 groups: training and testing to tune the hyper-parameters while accounting for overfitting and a validation group to approximate the accuracy per sample. Denovo peptides whose sequence was known with an accuracy of at least 90% were sequentially mapped against the Human proteome (HP) and a 3 Frame Translation (3FT) database. Since Leucine and Isoleucine are difficult to discriminate by MS, sequences with at most 1 leucine/isoleucine mismatch to any known protein were labeled “canonical peptides”. Similarly, peptides mapping to the 3FT database considered up to 1 leucine/isoleucine mismatch but were also stringently required to be at least 3 amino acids different from any known protein sequence before being considered non-canonical. **(b)** The resulting COD-dipp antigen library contains fields across the central dogma (genome, transcriptome, proteome). The COD-dipp web application allows for the development of physically templated neoantigens alongside additional statistics starting from public or patient-specific mutations to facilitate vaccine design.

#### Pipeline Components

To identify canonical peptides, MS-GF+ and Scavager^29^ were used as the closed search algorithm and to control the False Discovery Rate (FDR) to 1% respectively. To find and position post-translational modified peptides (PTMs), MSFragger and PTMiner^30^ were used to perform an open search analysis and to control both FDR along with False Localization Rate (FLR) to 1%. To find peptides from non-canonical sources, DeepNovoV2 was used for the *de novo* strategy (**methods**) in combination with the closed search PSM level information to train a specific DeepNovoV2 model per sample on interpreting the raw data. The training step for such deep learning approach is crucial for learning features of tandem mass spectra, fragment ions, and leverage sequence patterns in the immunopeptidome to impute over missing MS2 fragments. All high quality *de novo* peptides (90% accuracy) were sequentially mapped^31^ to the human reference proteome and afterwards to a 3 Frame Translation (3FT) database derived from the coding strand for each gene in the genome *i*.*e*., unspliced transcriptome (**cf. methods**). 3FT was used for the detection of peptides from novel sources (*i*.*e*., non-canonical peptides) such as introns, 5’UTRs, 3’UTRs, out of frame exons, and junctions spanning any of the previously mentioned features.

### A deep interrogation of immunopeptidomic datasets

Applying the COD-dipp pipeline across the dataset improved the number of peptide spectrum matches over any one strategy (**Fig. 3a**), revealing a breadth of canonical, post-translationally modified, and non-canonical peptides.

**Figure 3:**
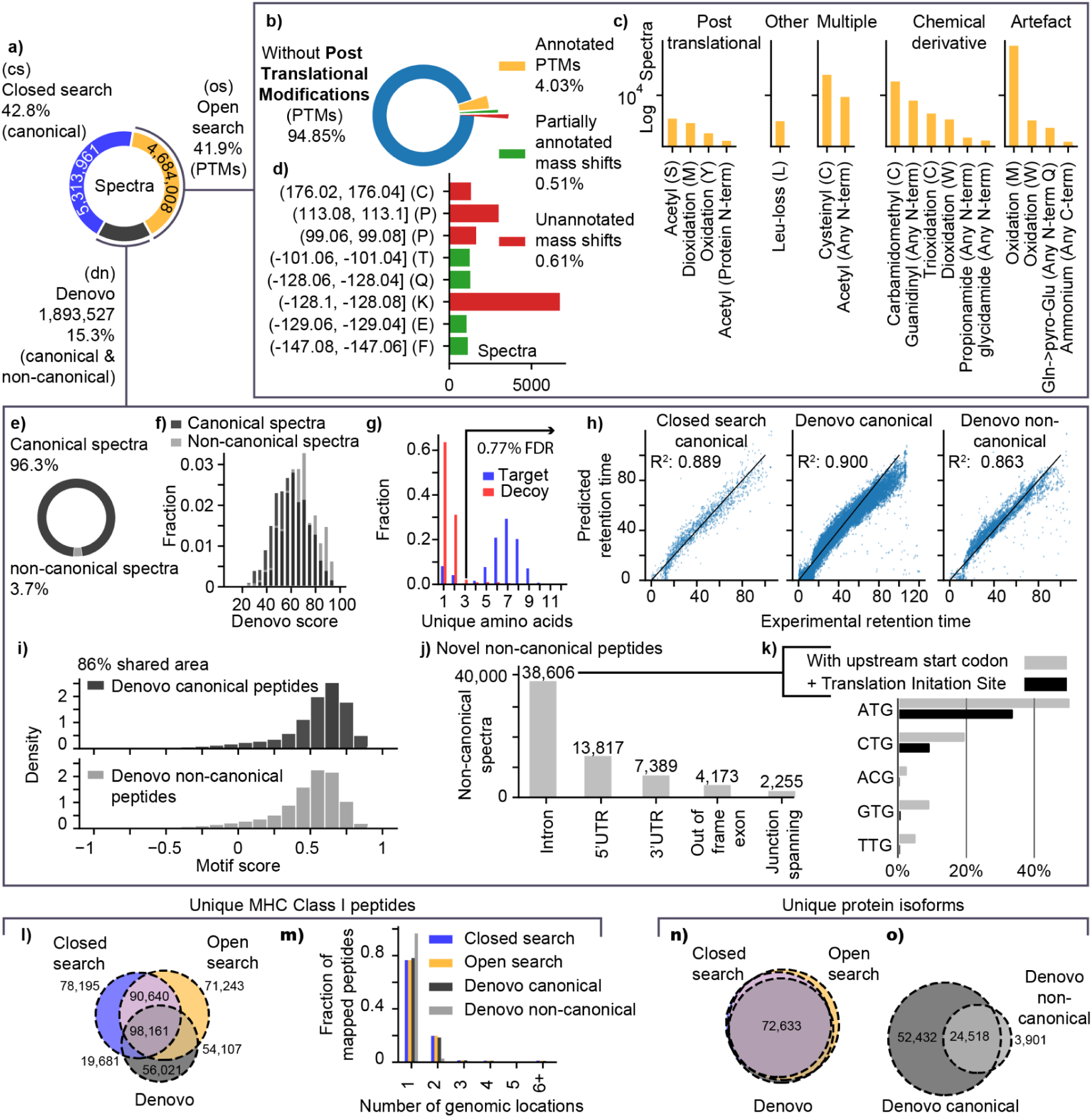
The revealed landscape of canonical, post-translationally modified and non-canonical peptide antigens. **(a)** Mass spectrometry peptide-spectrum match count per strategy. **(b)** Overview of Post-Translational Modifications identified by open search. (blue: spectra without PTMs, orange: spectra with a known UNIMOD PTM localized on a specific amino acid on the peptide. Green: The mass-shift is localized but the known PTM options do not fit the residue modified. Red: Otherwise. **(c)** Top annotated PTMs reported by modification type. **(d)** Most common partially annotated (green) or unannotated (red) mass shifts. **(e)** *de novo* identified spectra from canonical (dark gray) and non-canonical (light gray) sources. **(f)** Similar *de novo* score distribution of canonical and non-canonical spectra. **(g)** The use of target (blue) and decoy (red) frequencies to set a lower bound on the number of unique amino acids (minimum of 3) in an identified *de novo* peptide. **(h)** Correlation quality between predicted and experimental retention time for closed search and *de novo* peptides. **(i)** *de novo* peptides score similarly with motifs identified from database searches (Closed and open search) regardless of their canonical or non-canonical origin. **(j)** Distribution of non-canonical peptides into 5 categories of their origin. **(k)** Fraction of intronic peptides explained by the presence of an upstream start codon and predicted translation initiation sites. **(l)** Overlap of peptide identifications between strategies. **(m)** Fraction of peptides uniquely or multiply mapped to the genome per strategy. **(n)** Overlap of proteins identified between strategies. **(o)** Zoom in on the protein isoforms overlap between the *de novo* spectra from canonical and non-canonical sources.

#### Peptides with mass-shifts: Post-Translational Modifications (PTMs)

The robust Bayesian statistical analysis used in PTMiner for open search PTMs^30^ controls both False Discovery and False Localization Rates (FDR and FLR). Overall, 4,684,008 PSMs were detected at a 1% FDR + FLR by this strategy (**Fig. 3a**) and a subset of 5.15% was found to show PTMs (**Fig. 3b**). Cysteine (cysteinyl, carbamidomethyl and trioxidation) and Methionine (oxidation and dioxidation) were the 2 most modified residues (**Fig. 3c**). Interestingly, Cysteine has been reported as under-represented^32,33^ in comparison to the proteome and has been treated as a technical bias in the database search parameters previously. Furthermore, 1.12% of open search MHC Associated Peptides (MAPs) showed unknown mass shifts illustrated in **Fig. 3d** as green and red. Eight of the most common unexplained mass shifts were introduced by non-specific cleavage when combined with open search (**Supplementary Note 3**) except (176.02, 176.04] Da on Cys which remains with no clear assignable PTM in existing databases^34^.

#### Out of frame and intronic neoantigens

Several studies have explored MAPs from non-coding regions of the genome^17,35,36^ and novel antigens have been proposed from (I) pseudogenes and lncRNAs^37^ (II) intragenic non-coding regions^17^ through intron retention events^35^ or non-canonical translation events^36^. We developed a novel sequential approach for the specific purpose of detecting non-coding antigens from annotated genes. Our workflow identified 11,710 unique MAPs within intragenic non-coding regions. We explored the landscape of non-canonical antigen presentation in cancer using a rigorous sequential *de novo* sequencing strategy that made use of the entire immunopeptidomics dataset independent of matched genomics data (**methods**). We ensured that only high-quality spectra were assigned by applying a 90% accuracy threshold on *de novo* prediction. Carefully, preference was given for the human proteome before matching the non-canonical sequence space. Peptides were mapped to known normal proteins first, then the remaining to a 3 frame translation database based on the coding pre-mRNA sequences in GRCh38 as shown in (**Fig. 2, methods**). We did not examine fully non-coding genomic regions in order to focus on the most likely candidate peptides in this analysis, despite evidence that these too are translated^37^. Our *de novo* strategy contributed to 15.9% (1,893,527) of all identified spectra (**Fig. 3a**), with 96.3% mapping fully to normal exonic sequences (*i*.*e*., canonical) and a minority (3.7%) mapping to the 3 Frame translation database (*i*.*e*., non-canonical) sequences (**Fig. 3e**).

We assessed the quality of the de novo sequences by examining their DeepNovoV2 score quality, liquid chromatography retention times conformity, and HLA binding motifs appropriateness. Both canonical and non-canonical peptides showed a similar *de novo* score distribution with a slight shift of non-canonical peptides toward higher scores (**Fig. 3f**). The filtering of peptides with less than 3 unique amino acids *(i*.*e*., Poly A or poly G peptides) reduced the FDR (proportion of peptides that mapped to the decoy database) to 0.77% 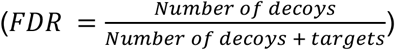 among exonic peptides (**Fig. 3g**). This filter was applied on non-canonical peptides as well. As another quality control step, the correlation between the experimental and predicted retention time of *de novo* canonical and non-canonical peptides was checked. **Fig. 3h** shows an r-squared score of 0.9 for *de novo* exonic and 0.863 for *de novo* non-canonical peptides in a melanoma sample (mel-15 of PXD004894) and an overall *de novo* non-canonical r-squared score of 0.88 among all samples. Similarly, *de novo* canonical and non-canonical peptides showed an 86% overlap in terms of HLA motifs indicating that the same MHC Class I haplotypes explain the newly found non-canonical peptides (**Fig. 3i, Supplementary Note 4**). All this provides strong evidence that these *de novo* peptides are high quality identifications (*i*.*e*., correctly predicted complete peptidic sequences). We found that any non-coding region type within genes can generate non-canonical peptides (**Fig. 3j**). **Fig. 3k** shows that most detected non-canonical peptides come from introns (58.3%) by either intronic retention or alternative Translation Initiation Sites (aTIS). Interestingly, we found that 86.29% of the intronic peptides had an upstream start codon and 40.65% to have a potential upstream Translation initiation site (TIS)^38^ hinting at alternative translation as a possible source.

#### Integrated search results

Of all 468,048 unique peptides, 21% were detected in all 3 strategies, and 19.4% by both closed search and open search (**Fig. 3l**). Closed search showed the largest exclusive set of peptides (16.7%) compared to open search 15.2% and *de novo* 12%. All strategies showed a similar number of mapped locations to the human genome with mostly 1 and 2 reference locations (**Fig. 3m**). Of all 85,208 unique protein isoforms, 85.2% (72,633) were detected by all 3 strategies (**Fig. 3n**). The low fraction of exclusive proteins reflects the complementary nature of the strategies. Similarly, the majority of non-canonical peptides identified by *de novo* sequencing originated from the same set of proteins of canonical peptides with 86.3% (24,518) of proteins overlap (**Fig. 3o**). This implies that non-canonical peptides originate from the same subset of proteins of canonical ones.

### Recurrent neoantigens from alternative sources

We further surveyed the recurrence of non-canonical peptides and their degree of ubiquity across cancer immunopeptidomes. 239 MAPs were recurrently identified in at least 10 samples with at least 2 spectra (**Fig. 4**). Some recurrent peptides occur exclusively within cancer types (colon cancer, melanoma, and ovarian carcinoma) while others are shared across cancer types. Reassuringly, 90.6% were predicted binders by NetMHCPan 4.0 to the HLA class I supertypes, 12 HLA alleles with binding properties that cover much of the human population. Strong binders were in the majority 76% (181), while 14.6% (35) were predicted as weak binders. In comparison, a set of 239 random peptides with random length from 8 to 12 shows only 5.7% strong binders, and 11.7% weak binders to the same 12 HLA types. In addition, these peptides were detected in colon cancer tissue without the help of HLA amplification treatments (IFN or TRAM) hinting at a high expression by the MHC class I system. 20 out of 239 peptides were found downstream of known frameshift mutations in COSMIC and were associated with low NMD efficacy scores^39^, which could offer an explanation for their origins. 87 out of 239 were exclusively shared between cancer samples without any occurrence in disease free samples of which 7 were found in COSMIC as frameshifts. In addition, the presence of some peptides could be explained by cancer aberrations affecting splice sites.

**Figure 4:**
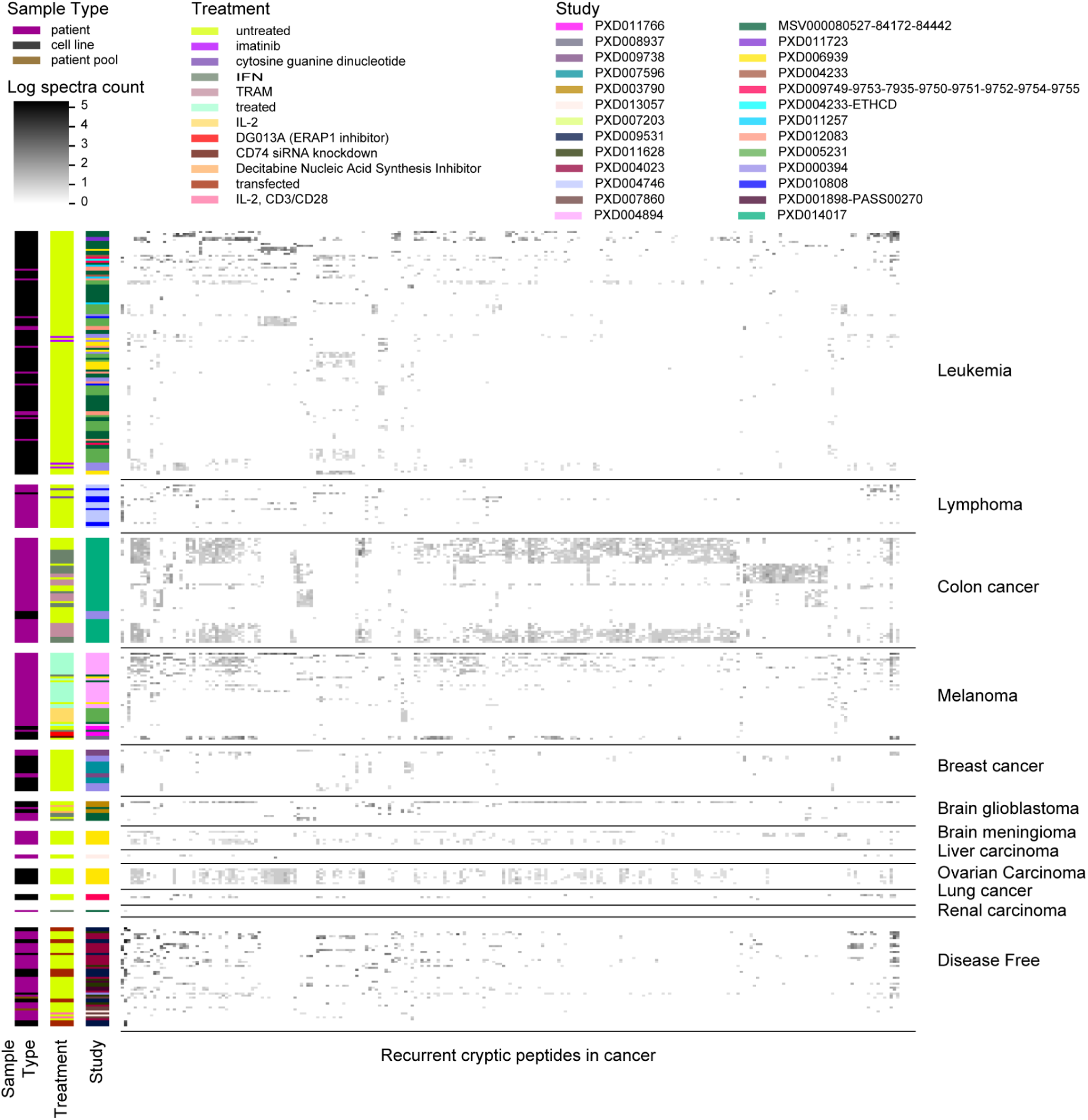
Recurrent non-canonical peptides in cancer. Heatmap showing the extent of shared non-canonical peptides (from *de novo* sequencing) across samples grouped by cancer type. Recurrent peptides were defined as sequences detected at least 2 times per sample and in at least 10 samples. In total 239 peptides from non-canonical sources passed this threshold. Each vertical row represents a recurrent non-canonical peptide and the gray to black intensity reflects the log10 spectra count for each peptide.

### Prognostic power of the COD-dipp generated antigen library

The assignment of somatic mutations to neoantigens tends to make use of predictive models of antigen presentation. Genomic mutations are translated to mutated sequences, then MHC Class I binding is evaluated for patient-specific HLA types using a sliding window to choose the best nonamer candidates. Here we suggest that neoantigen prediction should be informed by experimentally detected antigens (**methods**) and show a significant improvement over relying on predicted epitopes alone for immuno-therapy response prediction. **Fig. 5a** shows the performance of the homogenous fitness model by Łuksza et al.^40^ (top row) when using a typical prediction approach compared to focusing on genomic regions where antigens have previously been detected in our dataset (bottom row). The incorporation of mass spectrometry data improved the patient separation into responders (low fitness) and non-responders (high fitness) in all 3 cohorts. In addition, the 2 cohorts from Rizvi *et al*.^41^, and VanAllen *et al*.^42^ showed significance only with the mass spectrometry informed model.

**Figure 5:**
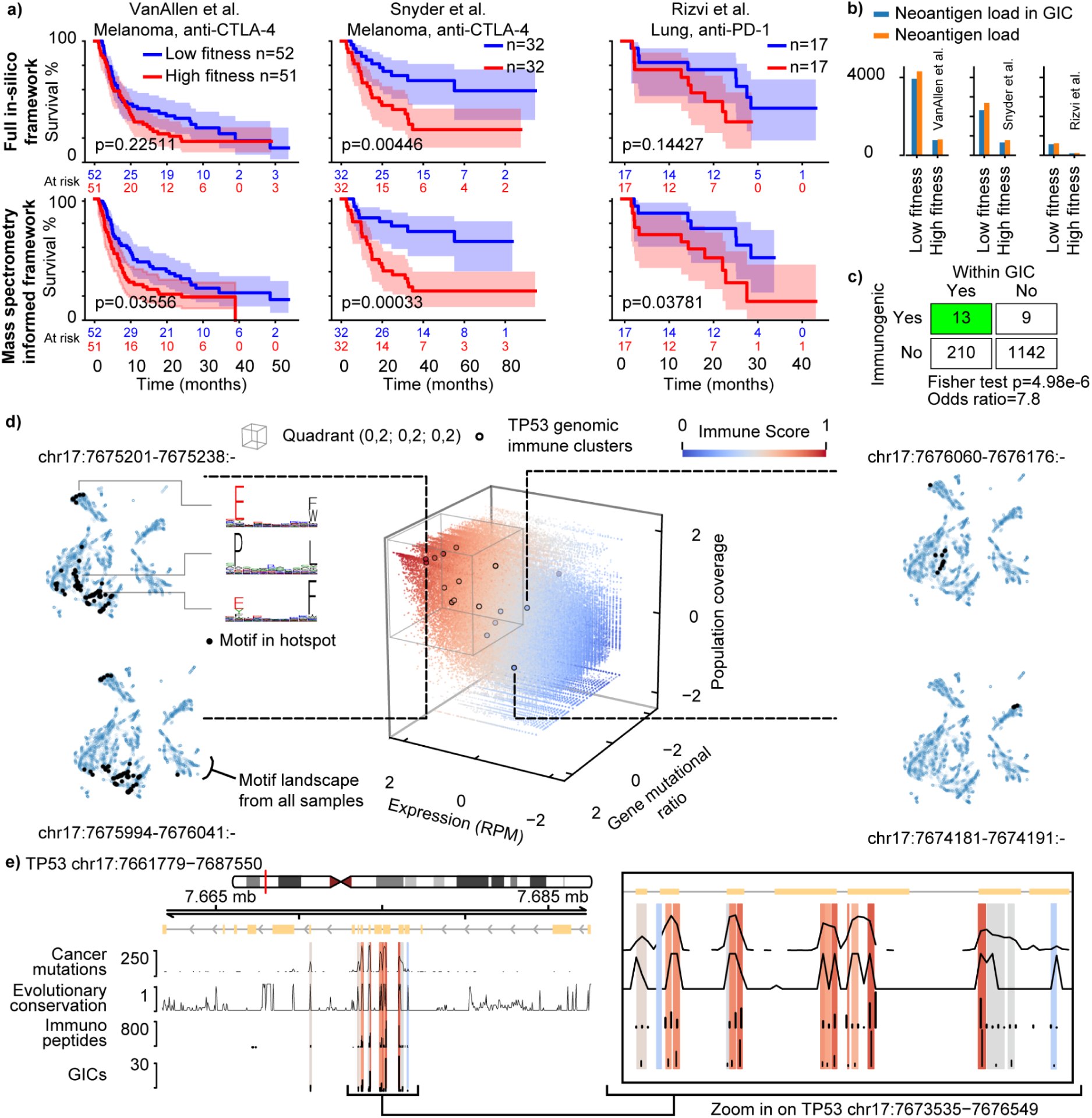
MHC Class I focal regions better predict immunogenicity and show diverse potential as vaccine targets. **(a)** Survival analysis of 3 cohorts using the “homogenous” fitness model from Łuksza *et al*. Patients are grouped into low fitness (immunotherapy responders) and high fitness (non-responders). Top row shows patient survival using only in-silico neoantigen prediction. Mutations in physically detected regions better predict overall survival (bottom row). **(b)** The majority of neoantigens (orange) lie in focal regions (blue) in low and high fitness groups. **(c)** a 7.8 fold enrichment in T-Cell reactivity within GICs versus outside GICs when using a combination of 1374 epitopes^43–45^. **(d)** The 140,966 focal regions of antigen presentation (Genomic immune clusters; GIC) differ in terms of antigen expression, mutational frequency in cancer (gene mutational ratio) and population coverage. As an example, focal regions in TP53 (black dots) have different properties in different regions. The 4 surrounding panels show the shared antigen presentation motif landscape across 486 samples (projected by UMAP). Each focal region presents peptides arising from different binding motifs and are therefore associated with different MHC-haplotypes so the breadth of coverage in motif space relates to population coverage of the immune-visible genomic region (**methods**) **(e)** Genomic coordinates for TP53 (on the left) highlighting immune clusters are according to the immune scores (IC) along with a zoom-in (right).

### Focal public neoantigens as simplified antigenic library

We define focal public neoantigens as sets of mutations from cancer-relevant hotspots that intersect with directly observed highly immune-visible regions in the genome. To find them, we first cataloged immune hotspots at the genome level. Although such hotspots have been reported before, here we paired them with a novel MHC haplotype deconvolution strategy **(methods, Supplementary Note 4; Supplementary Fig. S2)**. Hence, hotspots can be described in terms of their prevalence in the human immunopeptidome, across haplotypes and therefore in the population. All identified peptides from all samples were mapped to the human genome, and genomic coordinates from overlapping peptides were combined to define immune-visible genomic hotspots (Genomic Immune clusters or GIC, **methods, Supplementary Note 4**). **Fig. 5b** shows that the majority of neoantigens (orange) detected in the 3 previously mentioned checkpoint blockade immunotherapy cohorts are captured by focal regions (blue) in both low and high fitness groups. Furthermore, a combination of 1374 epitopes^43–45^ from studies evaluating the immunogenicity of cancer mutations revealed a significant 7.8 times **(Fig. 5c)** increase in T-Cell reactive epitopes within GIC regions (13/210) when compared to unreactive epitopes (9/1142) (two-sided fisher exact test p value: 4.98e-6). Antigen binding prediction by NetMHCpan 4.0 of these 1374 epitopes yielded only 4.8 fold enrichment in immunogenicity (two-sided fisher exact test p value: 3.34e-2). This indicates the value of verified immune-visible regions to identify neoantigens requiring 3 times less wet lab effort when pursuing T-Cell reactivity experiments. This simplifies the isolation of neoantigens, highlighting the importance of GICs for vaccine design. These genomic regions were associated with 3 features: (I) the mean MAPs expression calculated in Reads Per Million (RPM) (II) the cancer associated mutational density within the overlapping gene(s) (gene mutational ratio) (III) the percentage of the world population expressing the region given the deconvoluted HLA types **(methods, Supplementary Note 4)**. These features capture highly immune-visible genomic regions relevant to disease without a bias towards their width (**Supplementary Fig. S3**) and can be used to derive a score ranging between 0-1, which we call the *immune score* (**methods**). Finally, we developed an ‘immune score’ to rank focal regions according to the MHC Class I expression, the mutational load in cancer, and the population coverage according to the corresponding HLA types (**Fig. 5d; methods: immune score)**.

Interestingly, fourteen out of seventeen GICs of the tumor suppressor TP53 have a high *immune score* (**Fig. 5d, e**), indicating that multiple haplotypes converge on the same regions of this crucial cancer gene, which is consistent with immunogenicity profiles^46^ for TP53. Four examples in **Fig. 5d** illustrate how the diversity and spread of motifs in these genomic regions differ from region to region, and this relates to population coverage (**Supplementary Note 4 illustration**). High scoring GICs (**Fig. 5d** left side) are relatively more spread in the motif space in comparison to low scoring GICs (right side). This positive relation between the *immune score* and the population coverage is related to the cross-presentation of immune peptides across multiple HLA types (**Supplementary Note 4 illustration**). Interestingly, GICs with high *immune score* occur in regions of high evolutionary conservation according to the PHAST score^47^ (**Supplementary Fig. S4**), preference towards particular secondary structure elements **(Supplementary Fig. S5**), and are equally highly mutated in cancer. While 11.18% of the coding genome showed immune coverage (by at least 1 unique peptide), the GIC analysis revealed 140,966 regions of the human genome, that is 3.35% of the coding genome, being focal points of antigen presentation. Hence, the MHC system could be restricting presentation to focus on functional sites since they are mostly conserved throughout evolution.

### Feasibility of vaccines based on focal antigen presentation

We wanted to understand the feasibility of vaccine development using these focal regions of antigen presentation. Using the COSMIC^48^ database, pan-cancer aberrations were intersected with immune-visible regions of the genome and ranked by decreasing recurrence (**Fig. 6a**). The cumulative population penetrance (*i*.*e*., percentage of patients in COSMIC) increased to reach 38% when incorporating the top 30 mutations (**Fig. 6b**). Adding all mutations from these same immune-visible regions increased the population penetrance to 45% but required 2038 unique mutations. We noticed that certain cancer types tended to have recurrent mutations falling into the focal regions while others did not, indicating differences in cancer-specific potentials for vaccination. When the top 10 most recurrent mutations are considered, Hematopoietic neoplasms show a low vaccine potential, having low cumulative penetrance (11.9%) for immune-visible recurrent mutations. In comparison, large intestine (colon) carcinoma was highly immune-visible with 71.4% penetrance (**Fig. 6c**).

**Figure 6:**
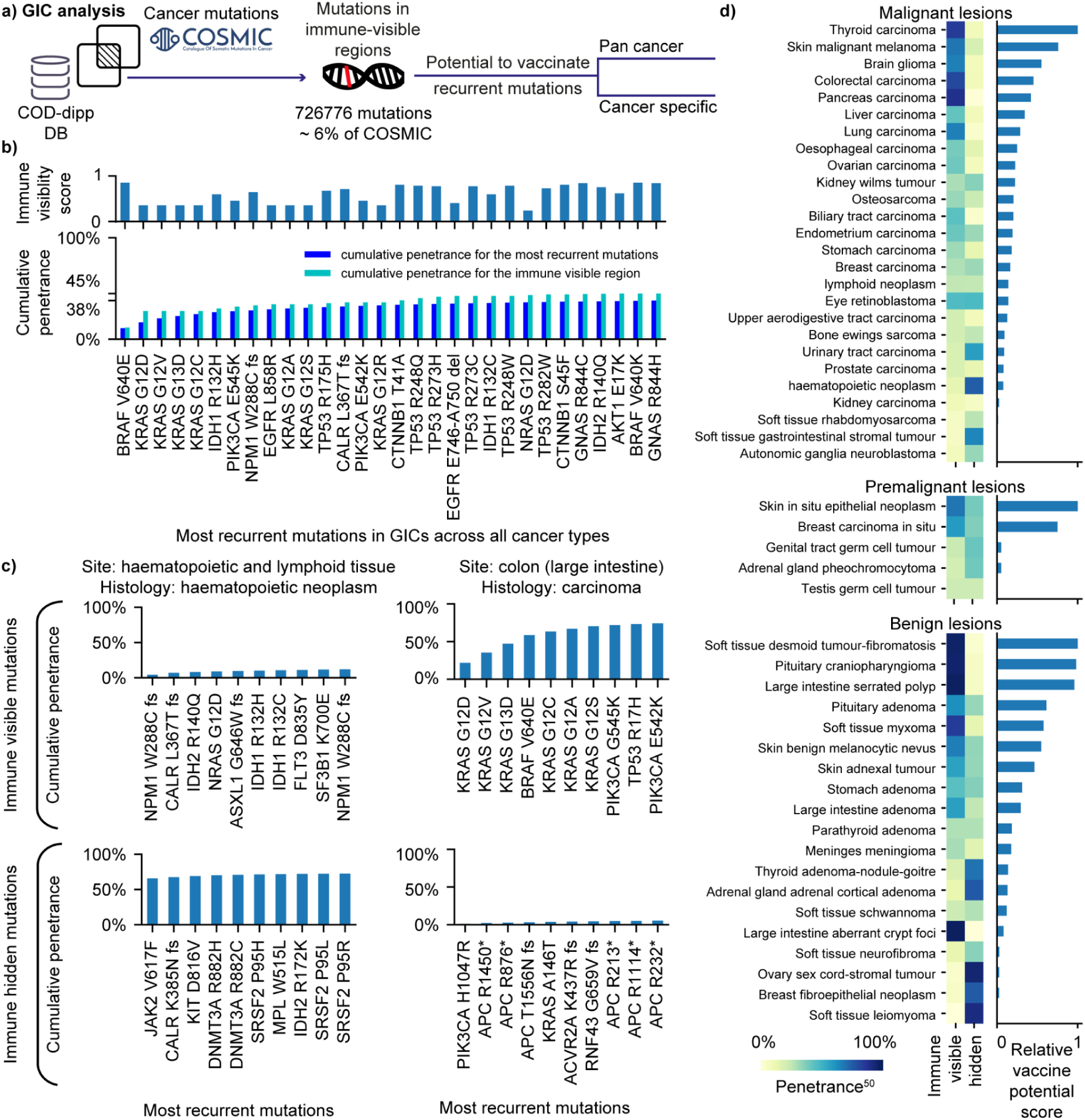
The tractability of building multi-epitope vaccines based on focal regions of antigen presentation. **(a)** Genomic immune cluster (GIC) analysis by intersecting focal regions (GICs) with cancer mutations from COSMIC database in a pan cancer (*cf*. panel b) and cancer specific fashion (*cf*. panels c and d). **(b)** Global analysis of the focal public neoantigen penetrance (proportion of people with a specific mutation in COSMIC) in all cancer types. Top bar plot shows the immune visibility scores for the 30 most recurrent aberrations in COSMIC. Bottom bar plot shows a cumulative penetrance calculated from the proportion of patients covered with each added mutation. (**c)** An example of a cancer with low vaccine potential (LHS: haematopoietic neoplasm) compared with one with high vaccine potential (RHS: colon carcinoma). A high vaccine potential is reflected by a high penetrance of immune-visible and low penetrance of immune hidden recurrent mutations. (**d)** Scope of vaccine potential across malignant, pre-malignant, and benign lesions. On the left, Immune-visible versus immune hidden mutational penetrance along with a relative score of vaccine potential on the right for each of visualized tumors found in COSMIC. Each cancer type was defined by the primary site and primary histology of the tissue.

We next developed a *vaccine potential* score (**Supplementary Note 5**) for a focal region that balances recurrence in cancer with additive penetrance. This *vaccine potential* score takes into consideration the proportion of people with a particular genetic mutation (penetrance) in a cancer type along with the immune-visibility on the MHC class I system generally across the population based on physically measured neoantigens in the current dataset. **Fig. 6d** illustrates the possibility to develop either therapeutic vaccines using public neoantigens across different premalignant and malignant lesions or vaccines against pre-cancerous or symptomatic benign tumors reported in COSMIC (**Supplementary Table 2**). The top malignant cancer candidates for potential vaccines are lung carcinoma, liver carcinoma, pancreas carcinoma, colon carcinoma, brain glioma, skin malignant melanoma, and thyroid carcinoma presented as the most attractive targets for therapeutic vaccines.

### A web application for vaccine development: understanding the immune-visibility of public and private neoantigens

To make the results presented in **Fig. 5** and **6** broadly accessible, we have developed a web application (https://www.proteogenomics.ca/COD-dipp) to facilitate vaccine design by bridging genomic mutations with physically detected MAPs (**Fig. 2b**). The portal provides a “neoantigen analysis” where the user can upload a set of gene mutations or a mutation calling file (VCF format) and retrieve their equivalent neoantigens templated from physically detected peptides. A second ‘GIC analysis’ feature is also included to give further information about the regions the mutations occur in. Mutations that overlap highly immune-visible regions, which we have called Genomic Immune Clusters (*i*.*e*., GICs) are returned. In addition, the expression levels, population coverage, enrichment in cancer mutations as well as the *immune score* of the immune-visible regions are provided (**methods**).

## Discussion

The cartography of antigen presentation developed by our open resource arises from a harmonized analysis of immunopeptidomics data mapped to the human genome. Our innovations over the most recent trends in computational mass-spectrometry identified a diversity of peptides mapping to the reference human proteome and its 3 frame translation. We mapped deviations away from the reference proteome as mass-shifts to reference peptides and explained significant numbers of these as genetic alterations (**Supplementary Note 6**) or PTMs. This approach expands with rigor on preliminary studies which either rely on proteogenomics for mutation calling^3^ or focus on the specific isolation and study of PTMs^18^. Our cartography is openly accessible as an alignment file directly usable by the genomics community to suggest focal neoantigens. On top of that, the easy access web-application, the high-throughput pipeline, and the code for all the analysis extends the accessibility.

### A diversity of peptides with post-translational modifications or from non-canonical sources

We found that 4% of all MAPs harbor PTMs. The most abundant detections are: carbamidomethylation, cysteinylation, oxidation, di or tri-oxidation, acetylation (**Fig. 3c**). Some PTMs are confirmatory chemical modifications from sample-preparation methods or common chemical derivatives (**Supplementary Note 3**). Other PTMs have been reported to increase immunogenicity of antigenic molecules against diseases^49^ and protect against degradation (**Supplementary Table 3**). For example, **Tri-oxidation** of cysteine has a potential of altering the immune response^50^, however its interaction mechanism with the HLA molecules and T cells is still in its infancy^51^. Additionally, T cells can discriminate **cysteinylated** from unmodified cysteine residues^51,52^. Likewise, N-terminal **serine acetylation** is known for multifunctional regulation, acting as a protein degradation signal, an inhibitor of endoplasmic reticulum (ER) translocation, and a mediator of protein complex formation. In our study, 96.3% of the cases of serine acetylation took place on P1 of peptides located at the second amino acid of proteins indicating the involvement of N-terminal acetyltransferases A after the initiator methionine is removed by methionine aminopeptidases^53^. P1 serine acetylation has been shown to protrude out of the HLA-peptide groove for T cell recognition^54^. With all the aforementioned, abundant PTMs implicated in immunogenicity such as serine N-term acetylation, cysteinylation, and tyrosine oxidation provide insights into immunogenicity and PTM-based vaccines.

The recurrence of non-canonical peptides within cancer types and between cancer types provides opportunities for vaccination. Several were found downstream of known frameshift mutations in COSMIC, which could offer an explanation for their origins and 87 out of 239 were cancer exclusive in our data. This includes 2 recurrent non-canonical peptides from an alternative exonic frame in the TSPO with an upstream splice variant (COSMIC ID: COSV61568369) that could cause a frameshift. The recurrence of certain non-canonical peptides in disease free samples is not surprising, since they contain normal tumor-adjacent tissue (21 samples) that could contain tumor contamination or PBMCs that might have been exposed to cancer cells and therefore do express neoantigen of non-canonical origins.

### Insights on vaccine and antibody-therapeutic design: Exploring the feasibility of recurrent and focal public neoantigen vaccines

Currently suggested biomarkers for response to checkpoint blockade immunotherapy include metrics based on tumor mutational burden and cell surface HLA expression. However, we now suggest a targeted exome-seq panel based on genomic hotspots of antigen presentation (The GICs in **Fig. 5d**) as a library to assess the likelihood of patients to respond to immunotherapy. Indeed, we have shown the prognostic value of templating neoantigens from previous physical detections **(Fig. 5a**) and their utility successfully shortlisting neoantigens (**Fig. 5c**). Indeed, we found that focusing on these regions can drastically simplify (7.8 fold) the identification of T-Cell epitopes. Likewise, this library may be valuable for informing both personalized and public neoantigens for vaccine development (**Fig. 6**) and we provide a simple web server to make this discovery accessible.

When recurrent mutations in COSMIC are intersected with focal hotspots of antigen presentation, the top 50 focal public neoantigens cover 78,326 patients in COSMIC (**Fig. 6a)**. The development of a multi-epitope vaccine requires further validation to confirm their presentation on more frequent HLA alleles as well as their immunogenicity. Vaccines covering more mutations within immune-hotspots could further broaden population coverage, but the number quickly rises to thousands of mutations with only small gains in sample coverage.

### Concluding remarks: An evolutionary perspective on focal antigen presentation

Our method and resource focus on making immune-surveillance accessible from the genome and transcriptome centric view, which may have far reaching implications that remain to be explored. It turns out that an analysis of recurrently identified peptides in the immunopeptidome reveals an enrichment of focal points in genes relevant to cancer. Of all the peptides detected in our study, 74% fell into focal regions that were conserved across vertebrate lineages (**Supplementary Fig. S4, S5**) and enriched in cancer genes (**Supplementary Fig. S3**). This would seem to indicate an evolutionary constraint on the immune system to preserve surveillance of these conserved regions.

Focused antigen presentation could have been converged on through two related processes. (I) fixing mutations in populations that maximize coverage of these regions by different haplotypes, or (II) by constraining the HLA molecules themselves to these regions. Indeed, genomic regions covered by more haplotypes may be constrained to code for anchor residues that allow broad presentation. Likewise, new HLA haplotypes may be constrained to maintain the presentation of these anchor regions. If the MHC Class I system tends to present focal regions important for cancer, then these regions could be prioritized for multi-epitope vaccines. Regardless of how, it would appear that evolution may have provided the right environment for the presence of focal regions in cancer genes on which to develop public and private neoantigen based therapies. These therapies could consist of multi-epitope vaccines spanning one or several regions that maximize coverage of the mutation landscape, either against one or multiple cancer types.

The tools and analyses herein may spark a new field of comparative immunology to understand how immune-surveillance changes due to the onset of diseases like cancer. Even between species a genome-centric view may help to better understand any evolutionary origins of genomic hotspots of antigen presentation. These fields would make use of genome and transcriptome centric AI models that can now be trained from our open and growing resource.

### Limitations of this study

The biological design of this study permits general claims to be made about neoplasms, whilst specific biological questions are out of scope. The findings in this paper may not be applicable for ethnic groups or regions of the world that were not included in the considered dataset. The mass spectrometry centric dataset limits our finding to the immunopeptidome fraction that falls into the dynamic range of the current technology. Most studies collected for analysis were based on immuno-precipitation and W6/32 antibody leading to a potential sampling bias toward W6/32 hla-types selectivity.

## Methods

### Dataset selection

Twenty-five studies were selected based on a list of keywords related to immunopeptidomics (**Supplementary Note 1**). Low-resolution analyses were eliminated and only MHC related datasets conducted with at least one of the following instruments Q Exactive, Q Exactive plus/HF/HFX, LTQ orbitrap velos, LTQ orbitrap elite, Orbitrap Fusion, Orbitrap Fusion Lumos were kept (**Supplementary Table 1**). An additional study^55^ was considered from the massive.ucsd.edu database as it incorporated 95 HLA-A, -B, -C and -G mono-allelic cell lines.

### Proteomic database generation

A protein database was downloaded using ENSEMBL (RRID:SCR_002344) r94 biomart, decoy sequences were appended by reversing the target ones and 116 contaminant proteins were added^56^.

As peptides with intronic and out-of-frame reading frames have been previously reported^17,36^, a pre-mRNA 3 frame translation database (3FTDB) was generated for protein coding genes based on ENSEMBL (RRID:SCR_002344) r94 using the AnnotationHub and biostrings R packages.

### Mass spectrometry computational analysis

The proprietary RAW files acquired from the instruments selected were converted to mzML and mgf format using MSConvert (ProteoWizard version 3.0.19295.c8b8b470d, RRID:SCR_012056) with the TPP compatibility and peakPicking filter on.

#### Database search strategies

PTM calling considered a 1% False Localization Rate (FLR) of mass shifts on peptides at specific amino acids (PTMiner) as well as a global False Discovery rate (FDR) of 1%. Similarly, closed search (for identifying canonical peptides) was restricted to 1% FDR using Scavager. MSFragger v2.2 search engine was used to conduct the open search analysis against the ENSEMBL (RRID:SCR_002344) r94 biomart protein database in combination with PTMiner v1.1.2 to apply a transfer False Discovery Rate (FDR) and a False Localization Rate of 1% (FLR, the rate of falsely localizing the site of modification). MS-GF+ v2019.04.18 (RRID:SCR_015646) was used for closed search against the protein database in combination with Scavager to apply an FDR of 1%. Both database search strategies considered 8 to 25 amino acid peptide lengths, unspecific cleavage and no fixed Post-Translational Modifications (PTMs).

#### *De novo* analysis

DeepNovo (v2)^19,20^ is a neural network based *de novo* peptide sequencing model that integrates Convolutional Neural Networks (CNNs) and Long short-term memory (LSTM) architectures to extract features from both the spectrum and the language of presented peptides. DeepNovo has demonstrated improved performance to the state-of-the-art *de novo* sequencing algorithms by large margins. The model can be tuned on a restricted peptide space to improve performance, and models were trained for each sample using spectra from closed search analysis. Validation and test sets were also derived from the closed search results. The trained models were used to perform *de novo* (predict) on the remaining unmatched spectra. *de novo* sequences with at least 90% accuracy were considered by thresholding the *de novo* prediction score considering the performance analysis on the test set.

#### *de novo* peptide annotation

*de novo* peptides coming from canonical human proteins were identified by a BLAT^31^ alignment against the protein target-decoy database. Sequences perfectly matching any protein sequence were considered exonic (1 mismatch allowed for isobaric amino acids Leucine and Isoleucine). All the remaining sequences unexplained by proteins were considered as potential non-canonical peptides and were aligned against the pre-mRNA 3 frame translation database. Stringently, peptides perfectly matching a 3 frame translation (3FT) sequence were required to have at least 3 mismatches with any known protein sequence before being considered non-canonical. Since PSMs can be assigned without complete sequencing accuracy, requiring a 3 amino acid difference alongside the 90% accuracy cutoff above, increases confidence that the peptides assigned fall far outside the standard human reference. Remaining *de novo* peptides without any canonical or non-canonical annotation were labeled as ‘unmapped peptides’ and discarded.

### Alignment of immunopeptides on the genome

Closed search, open search and *de novo* exonic spectra were converted to an mzTAB format and converted to proBAM format^57^ using an inhouse maintained fork of proBAMconvert^58^ to generate a proBAM format. *de novo* non-canonical spectra were converted to proBAM using the pysam python (RRID:SCR_001658) package^59^ according to the Proteomic Standard Initiative (PSI) specifications.

### Deconvolution of haplotypes

We aimed to characterize the landscape of focal neoantigens across tumors by overlapping the peptides discovered by our pipelines to the genome, and kept track of sample haplotypes in order to understand the population penetrance of each region. The compiled dataset was missing HLA-type information for 32.9% of samples and a further 52.5% were poly-allelic making complicated our understanding of the immunopeptidomes characterized and our discussion around focal points of antigen presentation. We focused on comparing HLA peptide binding motifs at the sample level in order to interpret and compare samples, and associated samples to haplotypes based on the binding motifs they contained. For each immunopeptidomics sample, we deconvolved haplotypes based on samples with known MHC haplotype (**Supplementary Fig. S2, Supplementary Note 4**).

We developed a method to visualize immunopeptidomics samples and to pool sets of sequences together representing ‘motifs’ related to the interaction interface between antigens and HLA molecules (**Supplementary Note 4**). UMAP projections of the immunopeptidomes (**Supplementary Fig. S2a**) revealed clear clusters of peptides in each sample that were inspected by generating Position Specific Weight Matrices (PSWMs), a commonly used representation of motifs (patterns) in biological sequences (**Supplementary Fig. S2b**). Clusters with at least 1 high and 1 mild conservation site were considered and labeled as high quality binding motifs. Thus for each sample, we were able to produce a set of confident ‘motifs’ denoted by a 20Xn vector representation of the PSWMs. In total we identified 6993 PSWMs across all samples (**Supplementary Table 4**).

We then developed a strategy to cluster, visualize and compare the motifs identified across all samples, which we dub the motif-binding landscape. To this end, we characterized the motif landscape by tracing similarities between all 6993 PSWMs from different samples using matalignerv4a to align matrices (**Supplementary Fig. S2c**). 248 Highly similar HLA Class I motif clusters were identified and covered 76.7% (5662) PSWMs.

Taking into account that 82.5% of the identified high quality motifs lacked HLA typing information, we developed a strategy to deconvolute HLA-types based on motif comparison. Hence, HLA type deconvolution was carried out by comparison against intra-cluster motifs coming from mono-allelic samples (**Supplementary Fig. S2d, Supplementary Note 4**). This imputation strategy increased the labeled motifs fraction to 85% (5943). The remaining 15% of motifs were either isolated in clusters without any mono allelic origin motif or were not assigned in clusters (**Supplementary Table 4**).

### Focal regions of antigen presentation

The detection of peptides overlapping a core genomic region (focal region) from patients with alternative HLA allotypes increases the presentation likelihood of mutations (neoantigens) in this region. Hence, peptides identified by open search, closed search and *de novo* (canonical + non-canonical) were aligned to the genome and pooled. Genomic immune clusters (GIC) were defined as overlapping peptides with a maximum distance of 24 nucleotides. 3 features per GIC were derived (1) Expression in reads per million (RPM): 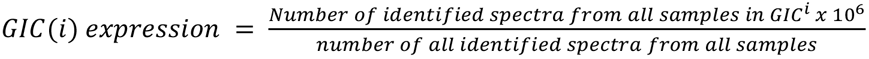 (2) population coverage: as a percentage of the world population that could be presented by considering at least 1 peptide from the cluster. This was calculated from the HLA types that the peptides in a specific GIC belong to^60^ (3) Gene mutational ratio (GMR): overlapping genes of each GIC were split into tiles of length 9 forming a set 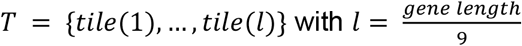. COSMIC mutations were counted in each tile then divided by the maximum count in *T* with 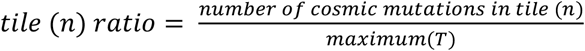. MGR(i) was defined as the maximum of t, a subset of *T*(t ⊂ T), consisting of tiles overlapping with the genomic immune cluster (i).

### Immune score

The 3 genomic immune cluster features (expression in RPM, population coverage, gene mutational ratio) were normalized using the powerQuantile method. An *immune score* was derived for each genomic immune cluster by multiplying the 3 normalized features producing an *immune score* ranging from 0 to 1. A high *immune score* reflects an increased MHC class I presentation, coverage of the world population, and relevance in cancer.

### Prognostics to immunotherapy response

Patients from 2 melanoma^42,61^ and 1 small cell lung cancer^41^ cohort were used to predict a response to immunotherapy. Genomic mutations from all patients overlapping the genomic coordinates of the mass spectrometry detected MHC peptides were kept. Neoantigens were generated by first introducing patient specific mutations to the WT genomic sequence of the MHC peptides followed by in-silico translation. NetMHCPan 4.0 was used to predict the binding of the wild type and mutated (neoantigen) immunopeptides using patients’ specific HLA Alleles. Both weak and strong binders NetMHCpan predictions were kept excluding all non binders. The Homogenous full AxR fitness model by Łuksza *et al*.^40^ was used for predicting response to immunotherapy. Survival plots and log rank tests were calculated using lifelines^62^ to compare the impact of a full in-silico versus mass spectrometry informed neoantigen framework on predicting response to immunotherapy.

### The pipeline architecture and technical details

Considering the complex nature of the immunopeptidomic database searches (unspecific cleavage) compared to proteomics (mostly tryptic cleavage) we implemented a pipeline with scalability in mind. The implementation is under snakemake v5.4.5, a pipeline manager, offering compatibility with most popular cluster workload managers such as SLURM. Therefore, a study with multiple patients would still take from 10 to 12 hours to complete on a cluster thanks to the parallel computations. In addition, the use of Conda, a package manager, allows the pipeline to automatically create software environments making it easily reproducible on other machines. For instance, the analysis of the pride dataset PXD004894 (*i*.*e*., 25 patients) comprised of 140 raw files took over 12 hours (real time) and around 28892 computational hours (∼5000 GPU hours for DeepNovoV2, ∼7000 CPU hours for MS-GF+, ∼16800 CPU hours for MSFragger, ∼92 CPU hours for Scavager).

## Code and data availability

1. The web portal: as an interface for easy neoantigen analysis and GIC analysis described earlier (https://www.proteogenomics.ca/COD-dipp).
2. The COD-dipp code: intended for High Performance Computing (HPC) will be made available as a snakemake pipeline on a git upon peer-review.
3. The full mass spectrometry processed library will be made available on figshare upon peer-review.

## Author Contributions

J.A and G.B. conceived of and initiated the project. J.A and S.K. coordinated and supervised the project. The first draft of the manuscript was written by G.B, J.A. G.B. collected the online studies, developed the computational approach and software, processed the data, made the figures and coordinated the manuscript. J.A and G.B developed the statistical methodology for the analysis. The manuscript was sent for revision and approval by all authors. A.L, C.B, F.M.Z, C. P., H.A, A.R, D. J. H, T.R.H, S.S part of the KATY consortium as well as T.W, M.Par, R.O, P.B and K.L. contributed to the writing of the manuscript. G.B, D.P, K.W., A.P, D.R.G, R.F., S.K, J. A. A. The International Centre for Cancer Vaccine Science contributed both the processing of the data and the writing of the manuscript.

## Acknowledgments

The International Centre for Cancer Vaccine Science project is carried out within the International Research Agendas programme of the Foundation for Polish Science co-financed by the European Union under the European Regional Development Fund. Authors would also like to thank the CI-TASK, Gdansk and the PL-Grid Infrastructure, Poland for providing their hardware and software resources.

