## Supplementary Notes for "The immunopeptidome from a genomic perspective: Establishing immune-relevant regions for cancer vaccine design"

**Note 1. Dataset selection** 2

**Note 2. Dataset processing** 3

**Note 3. Post translational modifications due to sample preparation or common chemical derivatives** 5

Illustration 6

**Note 4. Deconvolution of HLA haplotypes** 7

Peptide pairwise distance metric 7

Peptide Clustering to identify motifs 8

Clustering of motifs between samples and HLA-type inference 8

Denovo peptides motif similarity 9

Illustration 10

**Note 5. A score for vaccine potential** 11

Illustration 12

**Note 6. Single amino acid variants in the immunopeptidome** 13

**Supplementary Figure S1** 14

**Supplementary Figure S2** 15

**Supplementary Figure S3** 16

**Supplementary Figure S4** 17

**Supplementary Figure S5** 18

**References** 19

### Note 1. Dataset selection

**List of keywords used for selecting datasets from PRIDE:** Immunoprecipitation, Immunopeptidome, Peptidomics, Affinity purification, Mhc, Peptidome, Hla, Immunopeptidomics, Mhc class i, Ip, Hla peptidome, Hla-b*27, Hla class ii, Neoantigens, Immunoinformatics, Hla-c, Mhc class 1 ligands, Proteogenomic cryptic mhc lc-msms maps, Mhc class i antigen presentation pathway, Mhc-i peptides, Mhc i, Immunopeptidome; hla; lc-ms/ms; netmhcpan; binding prediction, Mhc ii, Mhc-i peptide-loading complex, Mhc affinity prediction, Mhc-ii peptidomics, Mhc ligandome, Mhc i-associated peptides, Mhc-i, Mhc class ii, Antigen presentation/ mhc class ii/ immunopeptidome/ peptide editing/ polymorphism, Mhc-i peptidomics, Shotgun proteomics; immunoprecipitation; meiosis; conserved proteins; meioc; , Anti-hla immunopurification, Immunopeptidome; hla; lc-ms/ms; netmhcpan; binding prediction, Personalized immunotherapy, Immunoprecipation, Immunoprecipiation, Immunoaffinity purification, Immunoprepicipitations, Immunopurification, Antigen presentation/ mhc class ii/ immunopeptidome/ peptide editing/ polymorphism, Hla-ii, Hla peptides, Hla-e, Hla-b*51, Hla class i peptides, Ducaf; hla-drb1*03:01, Hla typing, Hla-g, 'Hla class ı ligandome; hla class ı peptide ligands; high ph reversed phase; strong cation exchange; pre-fractionation', Hla-b40, Hla binding motifs, Hla-dm, Hla-b27, Immunopeptidome; hla; lc-ms/ms; netmhcpan; binding prediction, Hla-b*58:01, Hla-b*40:02 peptidome, Hla-dr peptides, Hla-dr, Hla-a, Hla-b57, Hla class i, Hla-i, Hla-a2, Hla-b, Interferon gamma; proteomic analysis; hla class i;  apm, Hla-i peptides, Hla-ligand, Hla-b*57:03, Hla-ligandomics, Hla-a*29:02, Hla-dr15, Hla-class i, Hla-restricted peptide

### Note 2. Dataset processing

We chose to work with Data-dependent acquisition (DDA) datasets, which were most abundant in the literature. In DDA, the instrument first cycles through a short MS survey scan ($MS^1$) of currently n eluting ionized peptides (10 or 20 most intense precursor masses) and a series of n (~10) MS/MS scans ($MS^2$)[1].

Computational methods are used to match the $MS^2$ spectra to an amino-acid sequence[2] in a process called peptide-spectrum matching (PSM)[3]. Closed-search, Open-Search and *de novo* sequencing are three main philosophies for carrying out this assignment and each one varies on the assumptions made about the contents of the sample. We chose one algorithm from each of these flavors of PSM assignment: MSFragger[4], MS-GF+[5] and deepNovov2[6] in order to broadly cover the spectra in all 486 samples.

Closed search (**Supplementary Fig. 1a**) is the most conservative strategy and assumes the peptides from the sample are derived from a specific set of reference protein sequences known as the search database. This set of sequences typically comprises a reference proteome, but can be augmented to include the various mutations identified through genomics and transcriptomics profiling. In closed search, an alignment attempt is made to identify peptides to the search database allowing a limited number of post-translational modifications, which must be specified a priori. In comparison, Open search (**Supplementary Fig. 1b**) is a less conservative strategy allowing the identification of peptides more distantly related to those in the search database. MSFragger[4] can identify post-translational modifications and single amino-acid variations (SAAVs) of the search database derived peptides by increasing the mass tolerance of the precursor ion tolerance. Contrasting these database search methods is *de novo* sequencing (**Supplementary Fig. 1c**), which assumes nothing about the sequences present in the sample and tries to derive them from a first-principles analysis of the $MS^2$ spectra[7–9].

Each strategy when applied alone suffers from a set of limitations. For instance, a closed search strategy[5] matches $MS^2$ spectra against peptides derived from the protein database and is the most widely adopted approach. However, it fails to detect even the most widespread modification known to exist on proteins, unless specifically included into the parameterization which leads to an exponential increase in computational time. An open search strategy[4] increases the amount of information that can be extracted by considering multiple post translational modifications (PTMs), by searching for peptides that exhibit a mass-shift away from the reference proteome. The source of the deviation can then be localized and attributed to mutations or post-translational modifications. It has not been widely adopted by the community due to two major bottlenecks previously unaddressed. Firstly, a reduced search speed that has been fixed by MSFragger[4], and secondly, a lack of algorithm for post-processing resolved not long ago[10–12]. Many samples lacked genomics data, so we present open-search results as a resource, but we note that there are significant caveats for mutant detection[13,14]. Likewise, *de novo* sequencing[6,7] which attempts to directly sequence the MS2 spectrum, hasn't been embraced due to lack of interpretability, quality control and post-processing algorithms. We addressed this issue by developing a strategy consisting of a sequential mapping to the reference, multiple filtering steps for accuracy control along with quality control checkpoints. These three philosophies of peptide-spectrum matching have never been combined in a single analysis data due to the lack of scalable pipelines and computational infrastructures. Used together, these 3 strategies deal with the remaining unexplained high quality MS/MS spectra. We have now developed a pipeline for the comprehensive characterization of antigens presented at the cell surface using this combination of algorithms.

### Note 3. Post translational modifications due to sample preparation or common chemical derivatives

Some PTMs are confirmatory, representing chemical modification from sample-preparation methods or are common chemical derivatives. For example, Cysteine **carbamidomethylation** is due to a reaction with iodoacetamide, used to block cysteine from oxidation. Nine out of the 26 studies used iodoacetamide in their lysis buffer. Interestingly, only one study (PXD006939) considered **carbamidomethylation** as a variable modification. This excludes a substantial fraction of cysteine containing peptides and should be addressed. Upon inclusion in our study we managed to recover 27,453 spectra matching 3,397 unique peptides to these modifications. Other PTMs like **Methionine (M) oxidation** (single most abundant PTM) and **dioxidation** (methionine sulfone) could be explained as chemical derivatives. Interestingly, methionine sulfone has been found to occur in-vivo in Proteus mirabilis[15], a gram negative bacteria present in malignant cancers[16], although it can result from the use of a strong oxidizing agent[17]. We also see PTMs that are extremely common on proteins such as **Protein N-terminal acetylation** which has different effects on proteins such as half-life time, folding properties and interactions.

Other mass shifts seen frequently remained unexplained after the open search annotation step. Particularly, a deviation of -128.1 to -128.08 Dalton on Lysine was detected over 6000 times. In most cases, it was located on the first 2 or last 2 amino acids within peptides (cf. illustration below). After further quality control, It proved to be an open-search identification introduced by non-specific cleavage. This was revealed by the open search post processing algorithm (PTMiner[10]) that we use. PTMiner, checks for mass shifts introduced by in-source fragmentation, nonspecific digestion or missed cleavages. It adds/deletes amino acids one by one (up to 5) from peptide N- or C-termini and checks the altered mass shift. Since Lys (K) mono isotopic mass is 128.09496 Da, open search assigns a sequence with an additional Lys and assigns it’s equivalent negative monoisotopic mass as a mass shift. Hence, when using open search in combination with non-specific cleavage (in the case of immunopeptidomics) one should correct for mass-shift introduced by non-specific digestion parameters. For this reason, these mass shifts should not be considered biologically relevant. The same applies for the unexplained mass shifts below:

- (113.08, 113.1] (P): non-specific digestion explained by an addition of I/L
- (99.06, 99.08] (P): non-specific digestion explained by an addition of V

- (-101.06, -101.04] (T): non-specific digestion explained by a loss of T
- (-128.06, -128.04] (Q): non-specific digestion explained by a loss of Q
- (-128.1, -128.08] (K): non-specific digestion explained by a loss of K
- (-129.06, -129.04] (E): non-specific digestion explained by a loss of E
- (-147.08, -147.06] (F): non-specific digestion explained by a loss of F

#### Illustration

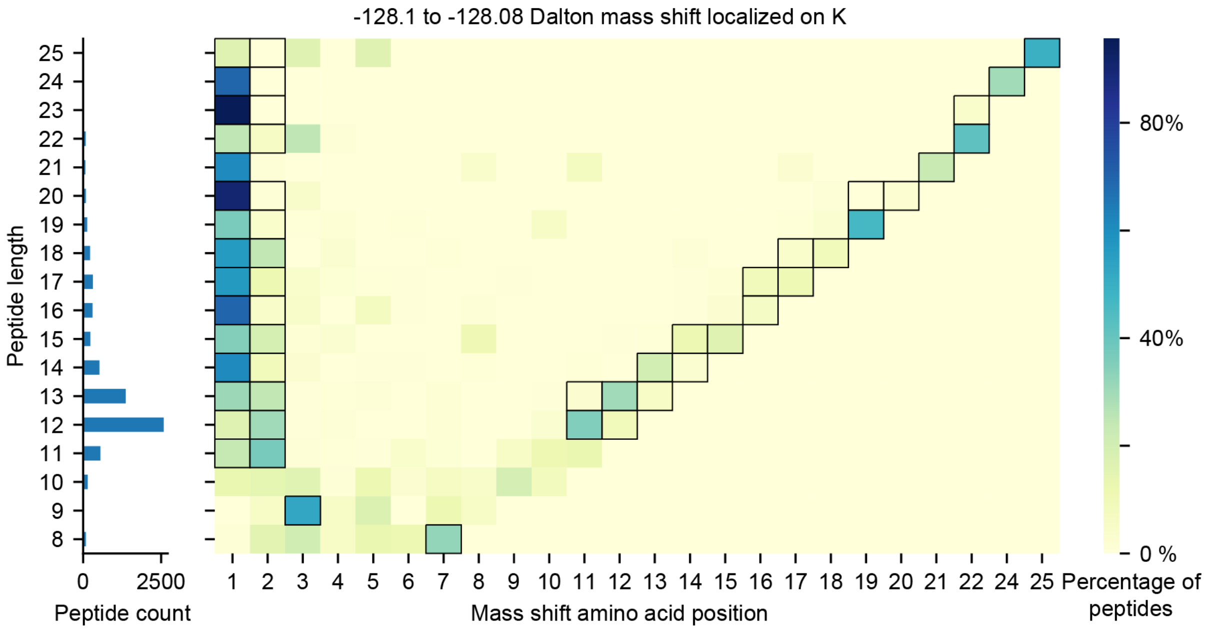


**Illustration:** -128.1 to -128.08 Dalton mass shift localization Illustration. Different peptide lengths are presented on the Y-axis versus the mass shift location within the peptides on the X-axis. The colors scale represents the percentage of the mass shift on each location within the peptide. The horizontal bar plot shows the number of peptides for different sequence lengths.

### Note 4. Deconvolution of HLA haplotypes

**Peptide pairwise distance metric**

We used a standard distance metric derived from phylogenetics to measure the distance between any two peptides. Assumptions of protein evolution models are supposed to closely match the evolutionary process to provide accurate phylogenetic estimates. The most widely used models of protein evolution assume evolution to be independent across sites and reversible according to a Markov model that relies on an instantaneous rate matrix **Q**. For any given alignment between two peptides, sites were treated independently and the likelihood of the exchange between the i'th and j'th amino-acid at each site was calculated as they would have been estimating exchanges across a branch in a phylogenetic tree: $p_{ij} = exp(Q_{ij} \times t)$ where t was set to unity (t = 1) reflecting the absence of a branch length.

**Q** can be decomposed into a set of stationary frequencies Pi_i for the model as well as a symmetric substitution matrix **S**. In short, $Q_{ij} = S_{ij} \times P_{ij}$. $P_{ji}$ represents the stationary or equilibrium amino acid frequency of each amino acid i that would arise at any site k if the Markov process were left evolving for a sufficiently long period of time. We chose the LG model[18] as the basis for our pairwise score function. The model specifies both S and $P_i$ and was downloaded from the authors' website (http://www.atgc-montpellier.fr/download/datasets/models/lg_LG.PAML.txt). We adjusted the stationary frequencies of the model to reflect the sample-specific amino-acid frequencies in the immuno-peptidome being studied. This adapts the standard evolutionary model to the patient-specific intricacies of the amino-acids presented by patient-specific haplotypes without having to know the MHC haplotype of the patient. Diagonal entries are obtained as the minus sum of the off-diagonals for the row and are thus proportional to the rate at which changes leave state *i*. To ensure that the interpretation of an edge length is the expected number of substitutions along that edge, **Q** is then rescaled so that $-\Sigma(P_{ii} \times Q_{ii}) = 1$.

When comparing two peptides, a sliding window approach was used to identify the linear alignment with highest probability arising by multiplying the likelihoods for each amino-acid exchanged between the aligned sequences. Peptides could technically share the same two anchor points but have different numbers of amino-acids between these anchors. Others have considered these peptides to share the same motif[19]. However, we have deliberately chosen to define these different spacings as different motifs by disallowing gaps. We think this is important, as they would necessarily fit differently into the binding groove, and these differences may prove valuable later in understanding immunogenicity. Pairwise scores were calculated between each pair of peptides to create a patient-specific scores matrix.

$$Score = Substitution\ score + (Substitution\ score - Best\ substitution\ score) \times Fraction \times Penalty$$

With $Substitution\ score = \Pi_1^k exp(Q_{ij})$, $Best\ substitution\ score = min(\Pi_1^k exp(Q_{ii}), \Pi_1^k exp(Q_{jj}))$ and $Fraction = \frac{Longer\ sequence\ length}{Shorter\ sequence\ length}$

where, k is the length of the shorter peptide, Q is the substitution matrix, and ij are the amino acids at a certain position of the first and second peptide respectively.

**Peptide Clustering to identify motifs**

A UMAP[20] manifold was applied separately to each sample pairwise matrix and clusters of peptides were labeled using Hierarchical Density-Based Spatial Clustering [21] (HDBSCAN). For each cluster, all peptides of the same length were grouped and all amino acid frequencies for all positions were computed. Then an euclidean distance per position was derived $E_i = \sqrt{\sum_{amino\ acids} (forground\ frequency_i - background\ frequency_i)^2}$ (*with* i being the position). At least 2 positions with euclidean distances greater than or equal to 0.5 and 0.3 were required to generate a position specific weight matrix (PSWM). Motifs were generated using logomaker version 0.8 [22] as a visualization of the PSWM matrices and scatter plots were generated using matplotlib [23].

**Clustering of motifs between samples and HLA-type inference**

Assessing sets of mutations from cancer-relevant hotspots (focal neoantigens) that intersect with highly immune-visible regions (public) in the genome requires association between antigens and HLA haplotypes. Taking into account that 82.5% of the identified high quality motifs within these regions lacked HLA typing information, motif comparison to deconvolute HLA-types was carried out by PSWM pairwise alignment using matalign-v4a[24], UMAP transformation and HDBSCAN clustering. Clusters consisted of highly similar motifs coming from different samples with known mono allelic HLA types (fully labeled), known poly allelic samples (semi-labeled) and unknown HLA types (unlabeled). HLA type deconvolution was done at the motif-level. Motifs coming from poly allelic samples were inferred by matching the closest mono allelic motif within the same cluster in case it fits the sample's HLA typing. HLA types of Unlabeled motifs coming from samples with missing HLA typing information were inferred by choosing the closest mono allelic motif within the same cluster.

**Denovo peptide motif similarity**

PSWM representations are useful to score the similarity between motifs (pattern) and any biological sequence (immunopeptides) having the same length as the matrix. Hence, we scored both denovo canonical and denovo non-canonical immunopeptides against the set of all motifs (6993 PSWMs from all 429 samples). **Fig. 3i** shows the normalized scores (the higher the score, the better the similarity) of denovo canonical and denovo non-canonical sequences against all motifs. Both canonical and non-canonical showed a skew towards higher scores reflecting a high similarity to HLA class I binding motif patterns. We already expected the denovo canonical peptides to show a high similarity to the set of HLA motifs knowing that they were used to generate the motifs themselves. However, a high similarity of denovo non-canonical peptides indicates a biological relevance/binding potential of these peptides to the HLA class I system. Moreover, denovo canonical and denovo non-canonical showed a 86% score distribution similarity (shared AUC) confirming, once again, that the identified non-canonical peptides are of high quality HLA binders.

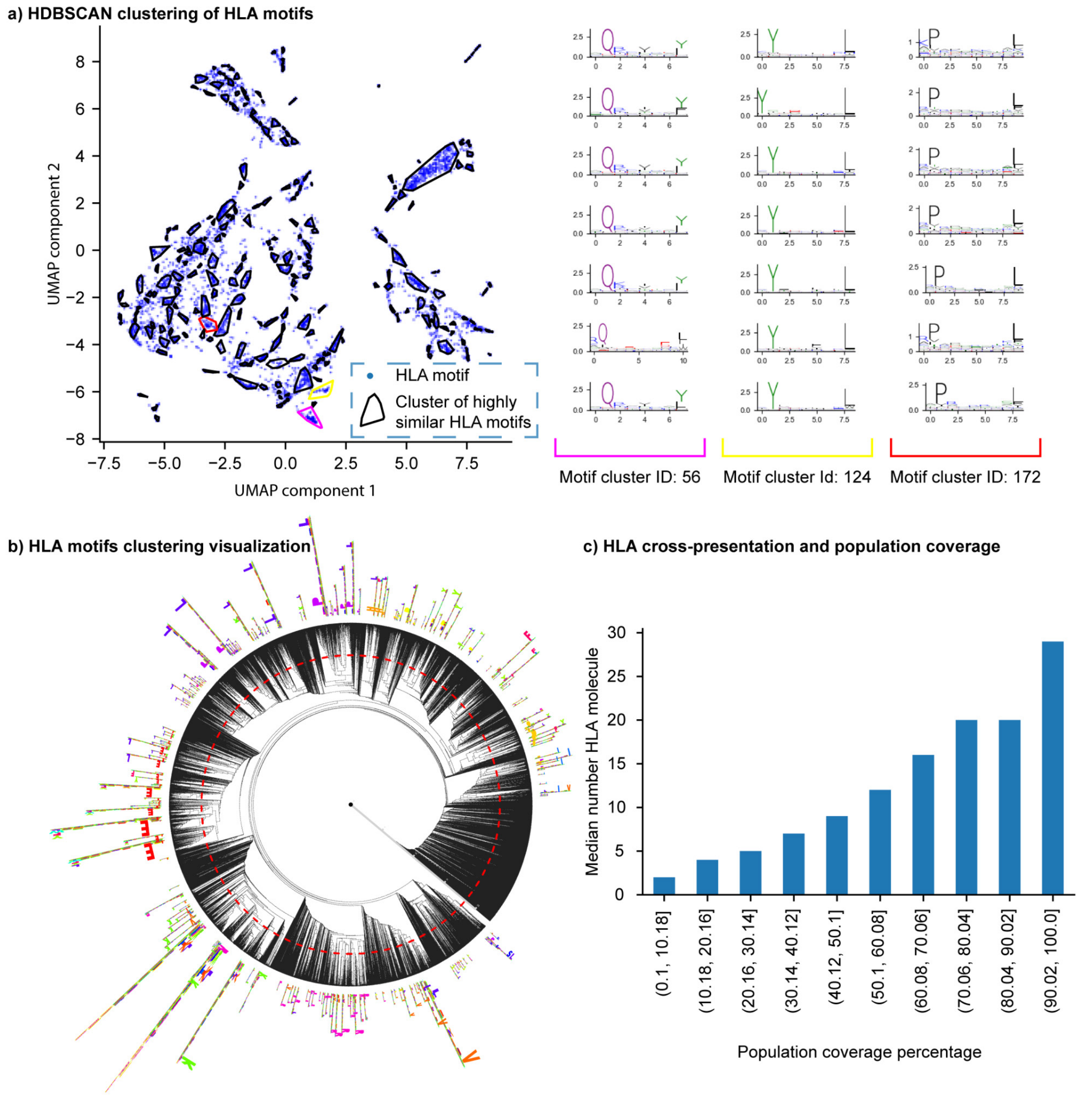


**Illustration: a)** Hierarchical Density-based Spatial Clustering of Applications with Noise (HDBSCAN) scatter plot after UMAP projection capturing the similarity between 6993 PSWMs (matrix representation of a motif) and assigning them into motif cluster represented by the contour lines. **b)** Motif clustering visualization with motifStack[25] showing the HLA anchor sites similarity. **c)** median number of HLA molecules for focal neoantigen regions (GICs) falling in the given population coverage percentage bins. As the population coverage increases the median number of HLA molecules per GIC increases as well. Hence, the triple correlation between MHC peptides expression, population coverage and number of HLA molecules per GIC infers that MHC expression and population coverage are correlated due to HLA cross-presentation.

### Note 5. A score for vaccine potential

We developed a *vaccine potential* score for a focal region that balances recurrence in cancer with additive penetrance $vaccine\ potential\ = \sum_{i}^{50} \quad (immune\ score\ x\ additive\ penetrance)^{i}$.

Here, penetrance is the proportion of patients with a particular genetic variant who belong to a certain cancer type, additive pentrance being the increase of penetrance with each added mutation and 'i' presenting the $i^{th}$ most recurrent mutation for a specific cancer type.

This vaccine potential score takes into consideration the penetrance of mutations in a cancer type along with the immune-visibility on the MHC class I system generally across the population based on physically measured neoantigens in immunopeptidomics studies detailed in the illustration below.

#### Illustration

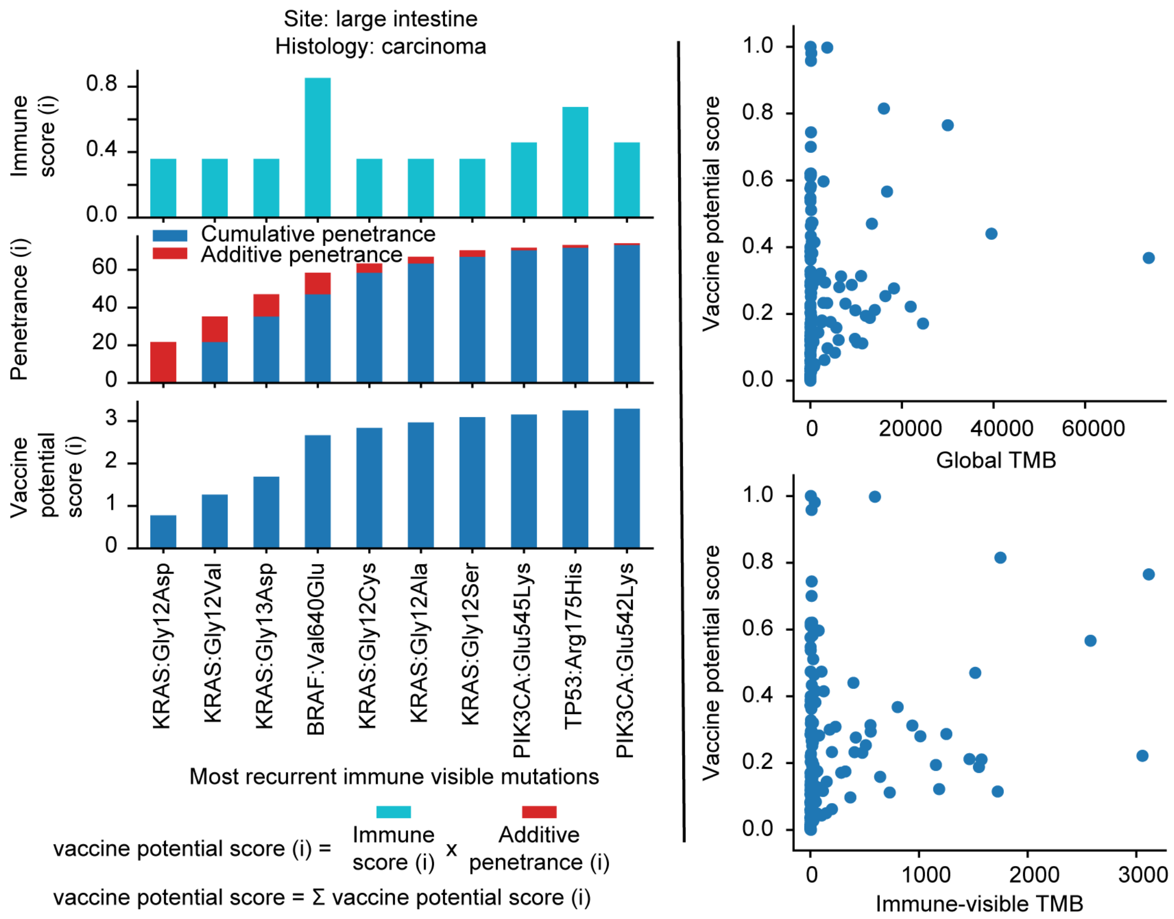


**Illustration:** schematic representation of the vaccine potential score on the left, taking into consideration the additive penetrance (red) and immune visibility score (cyan) of each added mutation to a multi-epitope vaccine. On the right, scatter of vaccine potential score versus global maximum TMB and Immune-visible TMB showing no relation between TMB and vaccine potential score.

### Note 6. Single amino acid variants in the immunopeptidome

Cancer is a disease of the genome, sculpted by the immune-system, which the tumor must avoid as it develops. However, the way these aberrations act on the immune-visible proteome is not trivial, and aberrations characterized by genomics can miss the heterogeneity of the tumor. For example, there can be post-transcriptional errors in mutation. Studies have demonstrated that the protein translation process from genetic material is prone to error. Misincorporation of amino acids during translation is estimated to occur once in every 5,000 codon on average[26,27]. These sorts of deviations, which would be absent from genomic evidence, elude typical MS-proteogenomics workflows that focus deeply on genomic evidence for mutation. In our analysis, since not all samples had matching genomics, we emphasized using open-search strategies to identify point mutations in reference proteins. The false discoveries related to using this methodology have been described[10], and we provide spectra for the mutations we have called. Our analysis has yielded multiple recurrent single amino acid variants across samples categorized as known polymorphisms, implicated in cancer or yet unreported. However, these variants lack support by exome sequencing due to the nature of our cohort and the broad view of focal neoantigens we aimed to analyze. Indeed, sequence coverage of typical MS closed search strategies is low[28] and known to miss mutations. On top of that, technical aspects may arise when considering a restricted search space of known proteins that doesn't exhaustively cover the MHC associated peptides. Hence, sequences originating from non-coding regions with a 1 amino acid difference with known proteins can mistakenly get categorized as SAAVs. Besides, the error-prone nature of mass spectrometry spectral matching can lead to false identifications.

In spite of the significant role of SAAVs in cancer and their prominent role in variant detection, their identification hasn't been very encouraging. For example, previous studies illustrate a recall of 0.49% of all non-silent genomic mutations in colorectal cancer[29] and 0.2% in melanoma[30] could be due to either a sensitivity limitation of the current technology[28]. Therefore, we prioritized discussing other sources of neoantigens in the main text by considering post-translationally-modified peptides, and novel open reading frames. However, the data for mutations is there and ready to explore.

### Supplementary Figure S1

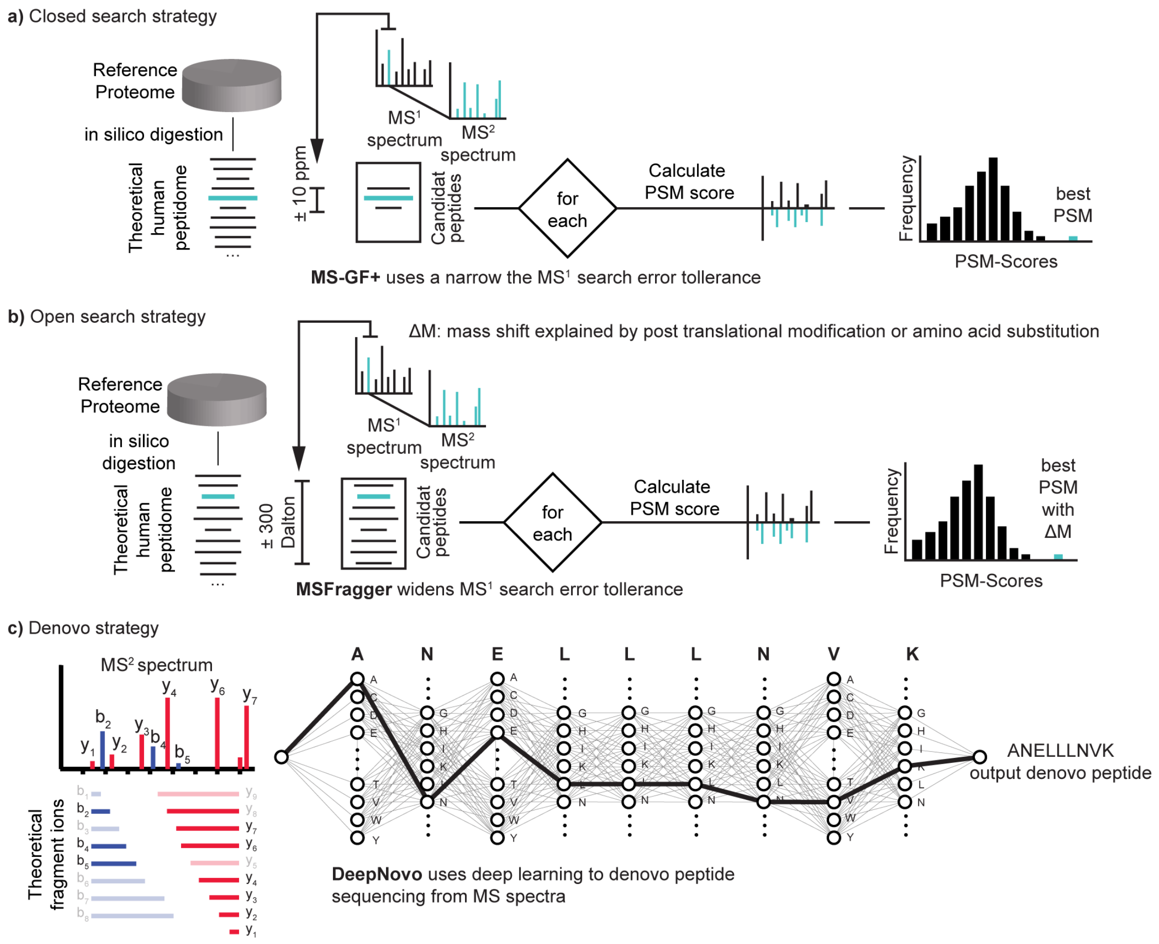


**Fig. S1: (a)** Closed search approach (supervised approach) requires a reference protein sequence database containing proteins expected within the sample. These protein databases are in silico digested and peptides falling within a certain error tolerance of the m/z window used to acquire the MS2 (the MS1 search error tolerance) are chosen as candidate peptide assignments. Each candidate peptide is then scored against the spectrum, using an algorithm-specific methodology and the candidate with the highest score is assigned as the sequence of the MS2 spectrum. **(b)** Open search strategy (semi-supervised approaches) widens the MS1 search error tolerance in order to identify peptides that would have been missed due to the mass-shifts caused by mutations and post-translational modification **(c)** *de novo* strategy (unsupervised approach) attempts to annotate spectra without a reference proteome by predicting a peptide sequence by directly reading the MS2 spectra.

### Supplementary Figure S2

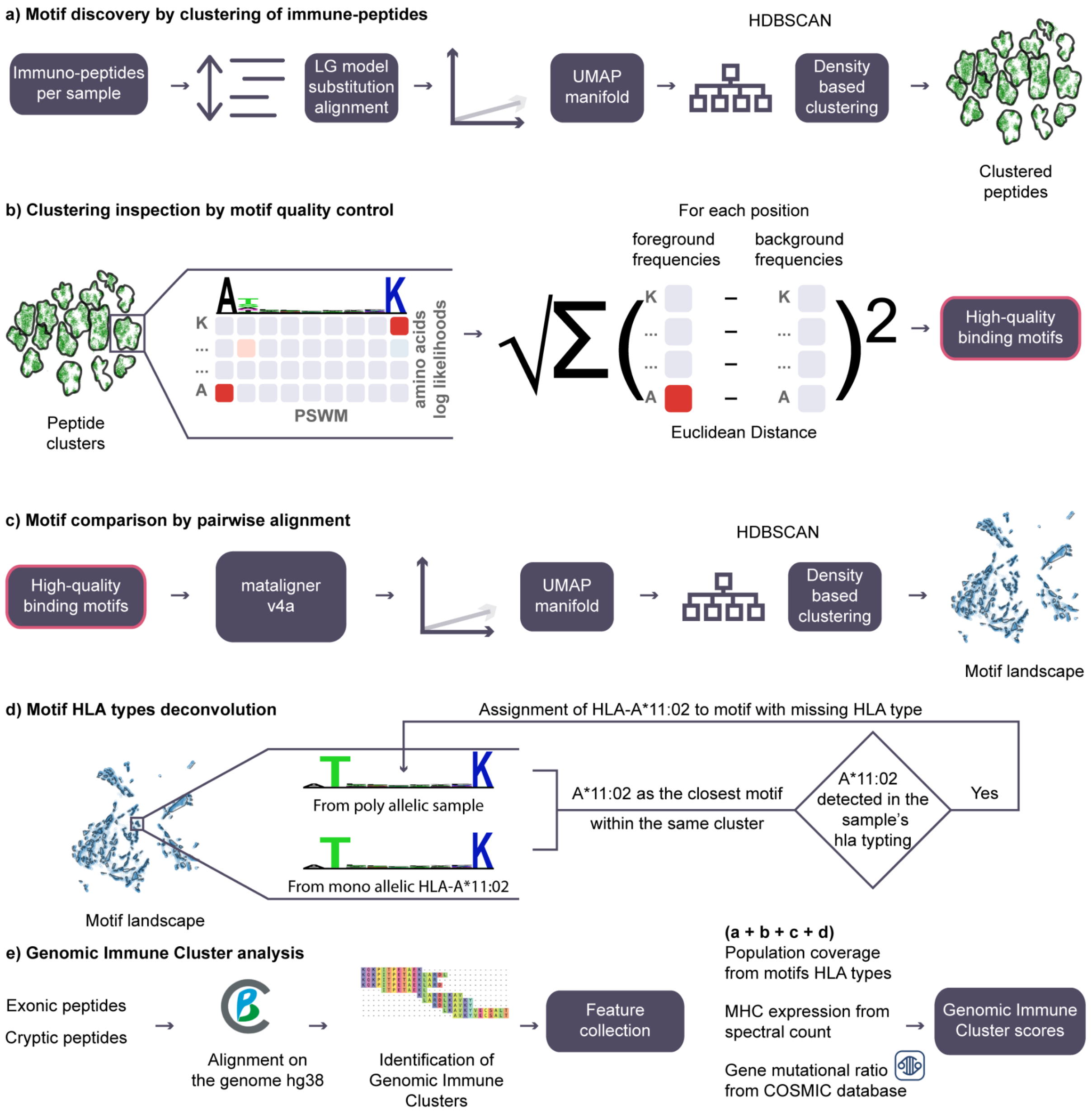


**Fig. S2: Computational methods schematic (a)** Sample-wise immune peptide clustering strategy using a pairwise comparison based on the LG substitution model **(b)** Quality control of resulting peptide clusters by motif inspection and filtering based on an euclidean distance metric. **(c)** Construction of intra-sample peptide binding motif landscape. Motifs were compared by Matalign-v4a and clustered by HDBSCAN to capture highly similar motifs . Similar motifs coming from different samples were developed by motif alignment. **(d)** Deconvolution of HLA types from motif data originating from polyallelic samples by comparison to mono allelic samples. **(e)** Identification of focal points of antigen presentation on the MHC class I system (genomic immune clusters: GICs) with a corresponding Immune Score (IC).

### Supplementary Figure S3

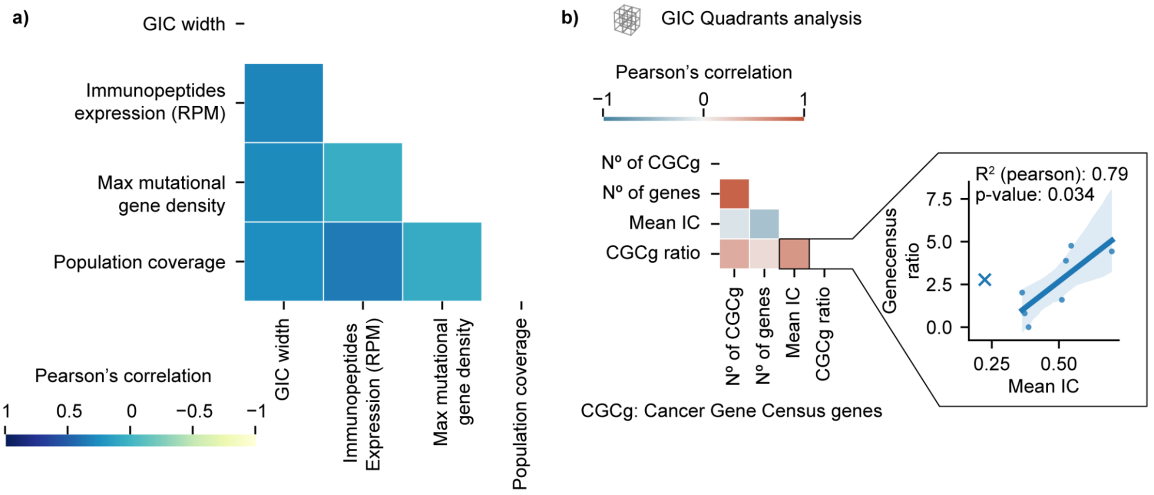


**Fig. S3:** Genomic immune cluster (GIC) features analysis. **(a)** Correlation plot between Genomic immune clusters (GICs) features and the width of the regions. This shows no correlation between any of Expression, Max mutational gene density and Population coverage with the region width. **(b)** Quadrant analysis (8 quadrants) regarding genes harboring mutations causally implicated in cancer (Cancer Gene Census genes or CGCg) and the Immune Score (IC). The heat map shows a pairwise correlation where GCGg ratio is defined as the number of CGCg / the number of genes. The relation between IC and CGCg ratio is depicted more accurately by the linear plot on the right. Quadrant (0,-2; 0,-2; 0,-2) shown as 'x' is considered an outlier restricting the analysis to an *immune score* range of 0.3 to 1.

### Supplementary Figure S4

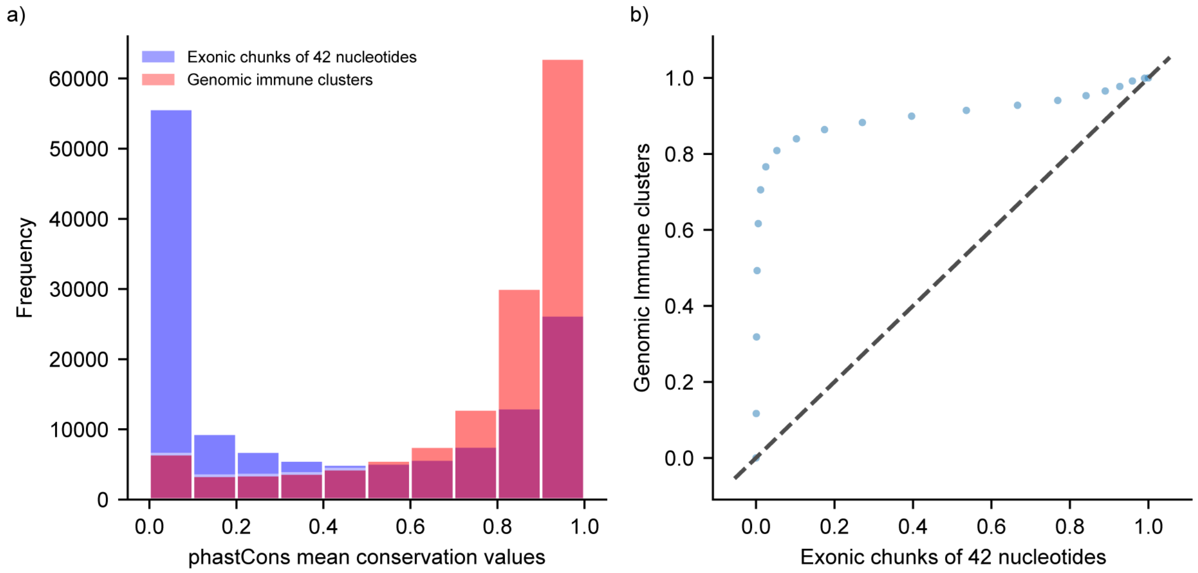


**Figure S4: Genomic immune clusters (GICs) mean conservation score (phastCons) versus non-covered Exonic regions split into chunks of 42 nucleotides (average GIC width). (a)** mean conservation score of GICs versus exonic regions split into chunks of 42 nucleotides showing a tendency of GICs to exist in conserved regions. **(b)** Q-Q plot showing two sets of quantiles against one another. If both sets of quantiles came from the same distribution, we should see the points forming a line that's roughly straight (dashed line). This plot shows a skew of GICs toward high mean conservation values (evolutionarily conserved regions).

### Supplementary Figure S5

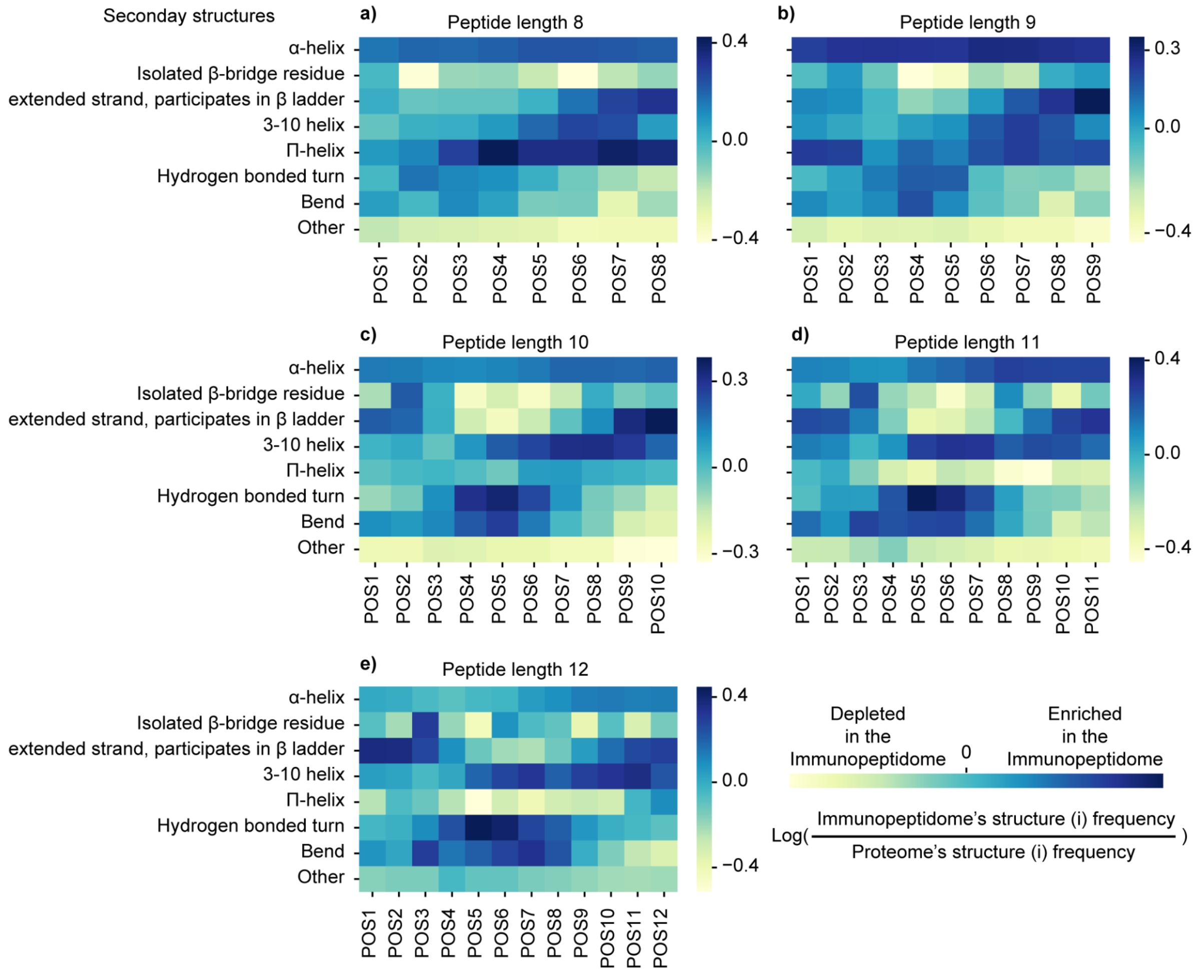


**Figure S5: Residue level analysis of the immunopeptidome's secondary structure composition**. Predicted structures of all human proteins by Alphafold2[31] were used to assign a residue level (amino acid) secondary structure using dssp[32,33]. A log ratio was derived reflecting the enrichment/depletion of secondary structures in the immunopeptidome in comparison to the proteome. Panels (**a** to **e**) present different peptide length groups (8 to 12 respectively). **(a)** length 9 peptides show a depletion of (Isolated β-bridge residue) on position 4 and 5, enrichment in Π-helix position 6 to 9, overall enrichment in α-helix and general depletion of the "other" structure. **(b-e)** show enrichment in 3-10 helix instead of α-helix (3-10 helix is more tightly wound, longer, and thinner than an α helix with the same number of residues). Perhaps, longer peptides tend to have longer helixes able to bend in the middle and bind. Also, the dominance of length nine peptides might be the result of their α-helix stability. In general, immuno-peptides seem to originate from structured regions of the proteome.

**Note**: the residues secondary structures here are relative to their location in proteins and do not reflect how they fold after proteasomal cleavage.
